# Quantitative Interactome Proteomics Reveals a Molecular Basis for ATF6-Dependent Regulation of a Destabilized Amyloidogenic Protein

**DOI:** 10.1101/381525

**Authors:** Lars Plate, Bibiana Rius, Bianca Nguyen, Joseph C. Genereux, Jeffery W. Kelly, R. Luke Wiseman

**Author notes:** These authors contributed equally to this work (Lars Plate, Bibiana Rius). To whom correspondences should be addressed: R. Luke Wiseman, Department of Molecular Medicine, The Scripps Research Institute, 10550 N. Torrey Pines Rd., MEM220, La Jolla, CA 92037.

## Abstract

Activation of the unfolded protein response (UPR)-associated transcription factor ATF6 has emerged as a promising strategy to selectively reduce the secretion and subsequent toxic aggregation of destabilized, amyloidogenic proteins implicated in diverse systemic amyloid diseases. However, the molecular mechanism by which ATF6 activation reduces the secretion of amyloidogenic proteins remains poorly defined. Here, we establish a quantitative interactomics platform with improved throughput and sensitivity to define how ATF6 activation selectively reduces secretion of a destabilized, amyloidogenic immunoglobulin light chain (LC) associated with Light Chain Amyloidosis (AL). We show that ATF6 activation increases the targeting of this destabilized LC to a select subset of pro-folding ER proteostasis factors that retains the amyloidogenic LC within the ER, preventing its secretion to downstream secretory environments. Our results define a molecular basis for the selective, ATF6-dependent reduction in destabilized LC secretion and highlight the advantage for targeting this endogenous UPR-associated transcription factor to reduce secretion of destabilized, amyloidogenic proteins implicated in AL and related systemic amyloid diseases.

## INTRODUCTION

The toxic extracellular aggregation of destabilized, amyloidogenic proteins is implicated in the onset and pathogenesis of diverse systemic amyloid diseases including Light Chain Amyloidosis (AL) and the transthyretin (TTR)-related amyloid diseases (1, 2). A critical determinant in dictating the pathologic protein aggregation central to these diseases is the aberrant secretion of destabilized, aggregation-prone proteins to the extracellular space (3). The efficient secretion of these proteins increases their extracellular populations available for concentration-dependent aggregation into toxic oligomers and amyloid fibrils that deposit in distal tissues such as the heart, inducing organ dysfunction. The importance of amyloidogenic protein secretion in disease pathogenesis suggests that targeting the biologic pathways responsible for regulating the secretion of destabilized, amyloidogenic proteins offers a unique opportunity to broadly ameliorate the pathologic extracellular protein aggregation implicated in the pathogenesis of diverse amyloid diseases (3).

Protein secretion through the secretory pathway is regulated by a process referred to as endoplasmic reticulum (ER) quality control (4–8. In this process, newly synthesized proteins are co-translationally imported into the ER where they interact with ER chaperones and folding factors. These interactions facilitate the folding of proteins into their native conformations and prevent their misfolding and/or aggregation within the ER. Once folded, these proteins are packaged into vesicles for trafficking to downstream secretory environments including the extracellular space. Proteins unable to attain a native, folded conformation within the ER are instead recognized by ER degradation factors and directed toward degradation pathways such as ER-associated degradation (ERAD). Through this partitioning between ER protein folding/trafficking and degradation pathways (i.e., ER quality control), cells prevent the secretion of destabilized, aggregation-prone proteins to downstream secretory environments.

In the context of systemic amyloid diseases, destabilized, amyloidogenic proteins escape ER quality control, allowing their efficient secretion to the extracellular space (1, 3). This suggests that enhancing ER quality control capacity could offer a unique opportunity to reduce the aberrant secretion and toxic extracellular aggregation associated with these disorders. One strategy to improve ER quality control for amyloidogenic proteins is by activating the unfolded protein response (UPR) (3, 9). The UPR regulates ER quality control through activation of UPR-associated transcription factors such as XBP1s and ATF6. These transcription factors induce overlapping, but distinct, subsets of ER chaperones, folding factors, and degradation factors (collectively ER proteostasis factors) that dictate ER quality control (10–13. The differential remodeling of ER quality control pathways afforded by XBP1s or ATF6 activation indicates that the independent activation of these pathways offers unique opportunities to correct pathologic defects in ER quality control for destabilized, amyloid disease-associated proteins (3).

Previous results show that stress-independent activation of XBP1s or ATF6 differentially influence ER quality control for destabilized amyloidogenic proteins such as ALLC – a destabilized Vλ6 immunoglobulin light chain (LC) associated with AL pathogenesis (14). Stress-independent XBP1s activation increases ALLC targeting to ER degradation pathways, while only modestly affecting its secretion (15). In contrast, ATF6 activation does not increase ALLC degradation, but significantly reduces the secretion and extracellular aggregation of ALLC. It does so without affecting secretion of an energetically normal Vλ6 LC or the endogenous secretory proteome (15, 16). ATF6 activation similarly reduces the secretion and toxic aggregation of destabilized variants of other aggregation-prone proteins, including TTR (10, 17–19). These results identify ATF6 as a potential therapeutic target that can be pharmacologically accessed to improve ER quality control and selectively reduce the secretion and subsequent aggregation of destabilized, amyloidogenic proteins implicated in amyloid disease pathogenesis (3).

Despite this potential, the molecular mechanism responsible for ATF6-dependent reductions in destabilized, amyloidogenic protein secretion remains poorly defined. Here, we establish an affinity-purification (AP)-mass spectrometry (MS) platform with improved sensitivity and throughput to define how ATF6 activation improves ER quality control to selectively reduce secretion of the destabilized, amyloidogenic ALLC. Our results define a mechanistic framework that explains the ATF6-dependent regulation of LC ER quality control and further motivates the development of therapeutic strategies that enhance ER quality control to ameliorate amyloid pathology in AL and related amyloid diseases.

## RESULTS & DISCUSSION

### Establishing an AP-MS platform to define ER proteostasis factors that interact with LCs

ER quality control processes are governed by interactions between non-native protein conformations and ER proteostasis factors (2, 6, 20). Thus, defining the molecular interactions between destabilized, amyloidogenic proteins and ER proteostasis factors allows identification of the components of specific biologic pathways responsible for dictating ER quality control for a given protein under defined conditions such as ATF6 activation. However, many challenges exist in defining interactions between ER proteostasis factors and destabilized protein substrates. These include the transient nature of substrate interactions with ER proteostasis factors and the difficulty in multiplexing interactome profiling to improve throughput without sacrificing sensitivity (21–26.

To address these challenges in the context of amyloidogenic LCs such as ALLC, we established an affinity-purification mass spectrometry (AP-MS) platform that utilizes Tandem Mass Tags (TMTs) for multiplexed quantification. We utilized this platform to define the specific ER proteostasis factors important for ATF6-dependent regulation of LC ER quality control (**Fig. 1A**). For these experiments, we employed HEK293^DAX^ cells (10), which exhibit ATF6-dependent reductions in the secretion of destabilized ALLC (14), but not the energetically-normal Vλ6 LC JTO (15, 27). We transiently transfected HEK293^DAX^ cells with flag-tagged ALLC (^FT^ALLC), flag-tagged JTO (^FT^JTO), or an untagged ALLC (**Fig. 1B** and **Fig. S1A**). These cells were then subjected to *in situ* crosslinking using the cell-permeable crosslinker dithiobis(succinimidyl propionate) (DSP) (28–30. We optimized DSP crosslinking to stabilize interactions between ER proteostasis factors and LCs in the ER (**Fig. S1B,C**). After crosslinking, we immunopurified (IP’d) ^FT^ALLC or ^FT^JTO using anti-Flag beads. Following stringent washing in high-detergent RIPA buffer to remove non-specific interactors, the samples were reduced to cleave the disulfide bond comprising the crosslinks, alkylated, and digested with trypsin. The digested peptides arising from individual experiments were then labeled with distinct TMT reagents, combined, and analyzed by Multi-dimensional Protein Identification Technology (MuDPIT) proteomics (31, 32). Specific recovery of peptides under different conditions was then quantified by comparing the recovered signals from the TMT reporter ions in the MS2 spectra (**Fig. 1A**).

**Figure 1.**
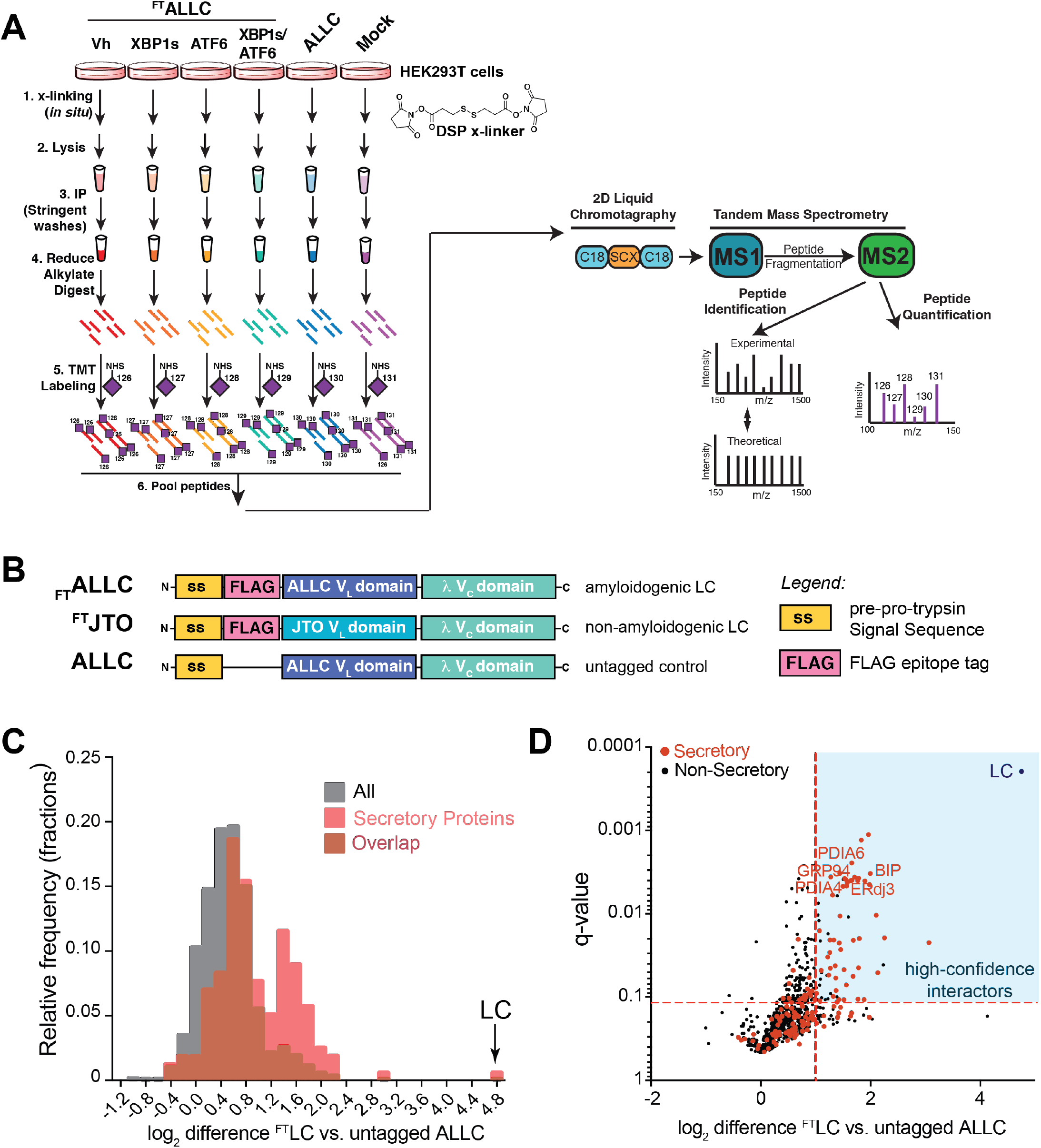
Establishing an AP-MS platform to identify ER proteostasis factors that interact with destabilized, amyloidogenic ALLC. **A**. Schematic of the multiplexed quantitative interactomics methodology, which combines affinity purification mass-spectroscopy (AP-MS) with in situ DSP cross-linking to capture transient, low affinity interactions with proteostasis network components. Sixplex tandem mass tags (TMT) are used for relative quantification of proteins in individual AP samples, followed by MuDPIT (2D LC coupled to Tandem mass spectrometry). **B**. Illustration showing the domain organization for the flag-tagged destabilized, amyloidogenic LC ALLC (^FT^ALLC), the flag-tagged energetically normal LC JTO (^FT^JTO), and untagged ALLC. A sequence alignment of ALLC and JTO showing the differences in amino acid sequence is shown in **Fig. S1A**. **C**. Histogram displaying TMT ratios of ^FT^LC (combined ^FT^ALLC or ^FT^JTO replicates) vs. untagged ALLC channels for all protein (grey) and filtered secretory protein (red). **D**. Plot showing TMT ratio (log_2_ difference ^FT^LC vs. untagged ALLC) vs. q-value (Storey) for proteins that co-purify with ^FT^LC (either ^FT^ALLC or ^FT^JTO) compared to untagged ALLC in anti-FLAG IPs. High confidence interactors are identified in the blue quadrant showing TMT ratio >2 and a q-value < 0.11. Secretory proteins are shown in red. Full data included in **Supplemental Table 1**.

Initially, we used this AP-MS platform to identify the ER quality control factors that bind LCs *in situ* by comparing TMT ratios for proteins that co-purify in anti-FLAG IPs from lysates prepared on HEK293^DAX^ cells expressing untagged ALLC or either ^FT^ALLC or ^FT^JTO (collectively ^FT^LC). We defined the TMT ratio as: TMT signal ^FT^LC IPs / TMT signal in untagged ALLC IPs. We observed two populations of proteins isolated in these samples separated by their TMT ratio (**Fig. 1C** and **Supplemental Table 1**). The first population exhibits a low TMT ratio of ~1.3, which represents proteins that non-specifically co-purify in both ^FT^LC and untagged ALLC anti-FLAG IPs. However, a second population of 72 proteins displayed a ratio of >2, indicating selective interaction with ^FT^ALLC and ^FT^JTO. This second population was enriched for secretory proteins and included ER proteostasis factors known to interact with LCs in the ER such as BiP, GRP94, ERdj3, HYOU1, and PDIA1 (33–42. We defined these 72 interacting proteins as *‘high confidence interactors’* of ^FT^LC (**Fig. 1D**), and we use these proteins as the basis for subsequent AP-MS experiments focused on defining the ER quality control pathways responsible for the selective, ATF6-dependent regulation of ALLC secretion.

### XBP1s or ATF6 activation differentially influence interactions between ER quality control factors and ^FT^ALLC

Stress-independent activation of XBP1s or ATF6 differentially influence ALLC ER quality control (15). XBP1s activation increases targeting of ALLC to degradation, while only modestly reducing ALLC secretion. In contrast, ATF6 activation significantly reduces ALLC secretion by 50%, but does not increase ALLC degradation, indicating that activating ATF6 increases the ER retention of this destabilized LC. In order to define the specific ER proteostasis factors responsible for the differential impact of ATF6 or XBP1s activation on ALLC ER quality control, we used our AP-MS proteomic platform to identify high confidence interactors that show altered interaction with ^FT^ALLC following stress-independent activation of these UPR-associated transcription factors in HEK293^DAX^ cells. These cells express both doxycycline (dox)-inducible XBP1s and a trimethoprim (TMP)-regulated DHFR.ATF6, allowing stress-independent XBP1s or ATF6 activation through the administration of dox or TMP, respectively (10).

We compared the recovery of TMT signals for high-confidence LC interactors that co-purify with ^FT^ALLC in lysates prepared from HEK293^DAX^ cells following 24 h treatment with vehicle, dox (activates XPB1s), or TMP (activates ATF6) (10). A challenge in comparing the TMT signals across different IPs is the variability of bait protein (e.g., ^FT^ALLC). ^FT^ALLC can vary in concentration owing to differences in transfection, variability in sample preparation, or alterations in protein secretion or degradation. Consistent with this, we observe 4-fold differences in ^FT^ALLC levels isolated from different replicates (**Fig. S2A**). This results in a large variance in unnormalized interaction ratios for high confidence interactors (**Fig. 2A**, blue). To address this variability, we normalized the recovery of high confidence interactors to the amount of ^FT^ALLC identified in each channel, significantly improving the variance across samples (**Fig. 2A**, orange) and allowing confident quantification of interactions changes. Importantly, alterations in the interactions between ER proteostasis factors and ^FT^ALLC observed using this normalization were nearly identical to those obtained using an alternative AP-MS approach that employed Stable-Incorporation of Amino Acids in Cell Culture (SILAC) for quantitation (**Fig. S2B-E**) (43–45. Comparing our multiplexed TMT-based platform to SILAC quantification also demonstrated other advantages of using TMT for these types of interactome studies. Since SILAC quantification only enables binary comparisons, a higher number of mass spectrometry runs and more instrument time was needed to generate the quantitative comparisons between different conditions (**Fig. S2F**). Furthermore, the number of proteins that could be reliably quantified in at least 3 biological replicates was at least 4-fold greater using our TMT-based platform than in any of the pairwise SILAC comparisons. (**Fig. S2F**). The improved throughput and increased sensitivity of our TMT AP-MS platform highlights the advantage of this strategy for defining interactome changes for destabilized proteins such as ^FT^ALLC induced by different conditions including XBP1s or ATF6 activation.

**Figure 2.**
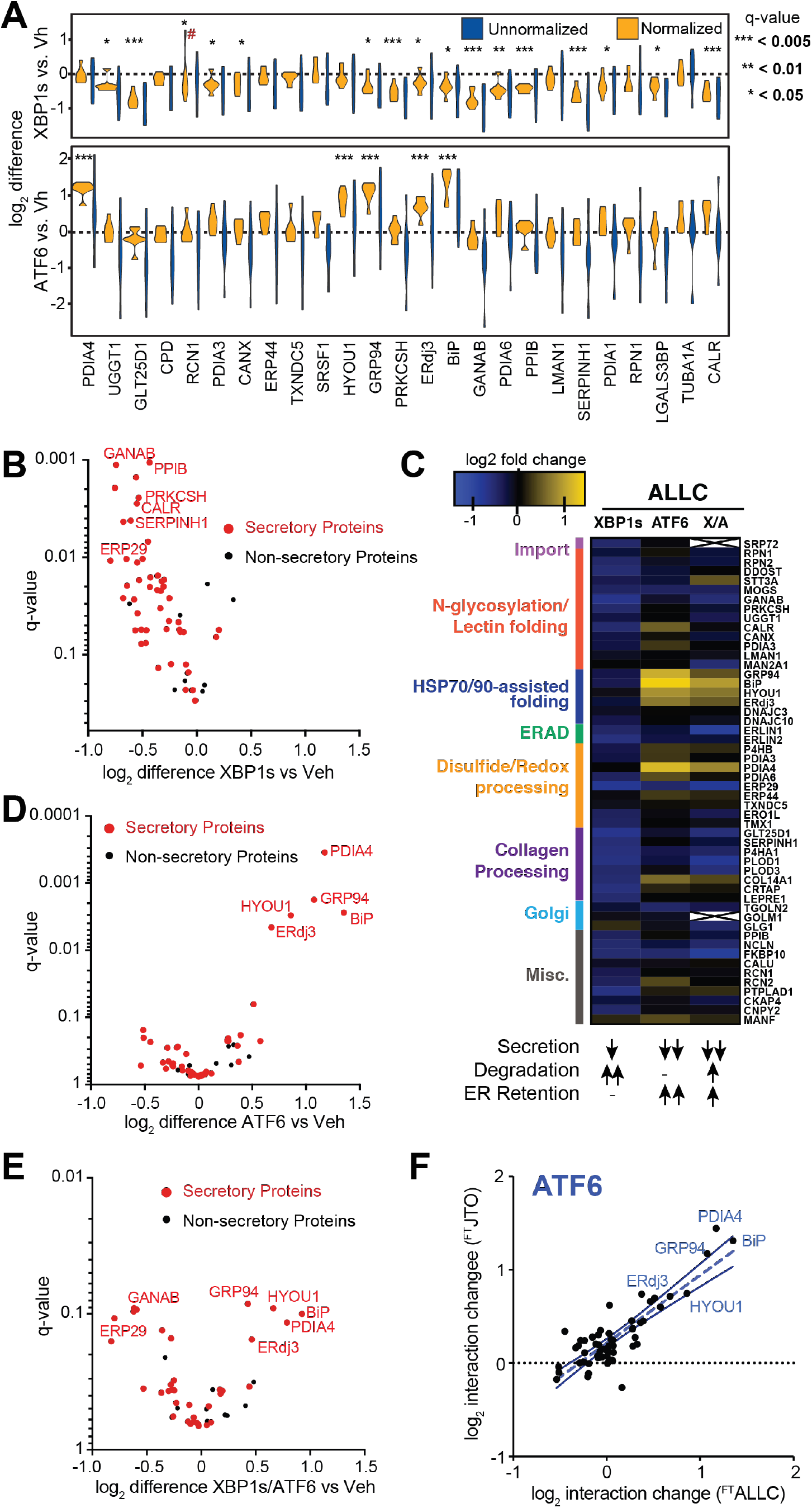
Stress-independent XBP1s or ATF6 activation differentially influence interactions between ^FT^ALLC and ER proteostasis factors. **A**. Plot showing the distribution of unnormalized (blue) and ^FT^ALLC-bait-normalized (orange) TMT interaction ratios for n = 6 – 7 biological replicates comparing the recovery of high confidence ALLC interacting proteins in anti-FLAG IPs from cells following XBP1s activation (top) or ATF6 activation (bottom). A simple normalization procedure of the protein TMT signal against the ^FT^ALLC bait protein signal across each TMT channel greatly diminishes the variance in interaction ratios. *q-value (Storey) < 0.15, **p < 0.05, ***p < 0.01; # denotes excluded outlier **B**. Plot showing TMT interaction ratio vs. q-value (Storey) for high confidence ^FT^ALLC interacting proteins that co-purify with ^FT^ALLC in HEK293^DAX^ cells following stress-independent XBP1s activation. Full data included in **Supplemental Table 2**. **C**. Heatmap displaying the observed interactions changes between ^FT^ALLC and high confidence ER proteostasis network components following stress-independent XBP1s or ATF6 activation. Interactors are organized by pathway or function. The previously defined impact of activating these pathways on ALLC secretion, degradation, and ER retention is shown below (15). **D**. Plot showing TMT interaction ratio vs. q-value (Storey) for high confidence ^FT^ALLC interacting proteins that co-purify with ^FT^ALLC in HEK293^DAX^ cells following stress-independent ATF6 activation. Full data included in **Supplemental Table 2**. **E**. Plot showing TMT interaction ratio vs. q-value (Storey) for high confidence ^FT^ALLC interacting proteins that co-purify with ^FT^ALLC in HEK293^DAX^ cells following stress-independent XBP1s and ATF6 coactivation. Full data included in **Supplemental Table 2**. **F**. Plot comparing the interaction changes of high confidence ALLC interacting proteins with either ^FT^ALLC or ^FT^JTO following stress-independent ATF6 activation. The dashed line represents least-squares linear regression. The solid lines show 95% confidence intervals

Interestingly, XBP1s or ATF6 activation induce distinct changes in the interactions between ^FT^ALLC and ER proteostasis factors, reflecting the distinct impact of these UPR-associated transcription factors on ALLC ER quality control (**Fig. 2B-D** and **Supplemental Table 2**)(15). XBP1s activation globally reduces interactions between ^FT^ALLC and ER proteostasis factors (**Fig. 2B,C**). This is consistent with the XBP1s-dependent increase in ALLC targeting to ER degradation pathways (15). Unfortunately, components of degradation pathways were poorly detected in our proteomics samples, which likely reflect these mainly membrane-associated proteins requiring specific detergents for solubilization (46).

In contrast, ATF6 activation increases interactions between ^FT^ALLC and select ER proteostasis factors, including the ATP-dependent ER chaperones BiP and GRP94, the BiP co-chaperones ERdj3 and HYOU1, and the protein-disulfide isomerase PDIA4 (**Fig. 2C,D**). The increase in ^FT^ALLC interactions with these ER proteostasis factors is consistent with the ATF6-dependent increase in ALLC ER retention (15) and suggests that ATF6 activation reduces secretion of ALLC through the increased targeting of this destabilized LC to specific ER proteostasis pathways.

We next sought to define how the combined activation of these transcription factors influences the ^FT^ALLC interactome. Despite impacting ALLC ER quality control through distinct mechanisms, co-activation of XBP1s and ATF6 does not synergistically influence destabilized ALLC secretion (15). Instead, XBP1s and ATF6 co-activation reduces ALLC secretion to the same extent observed with ATF6 activation alone and modestly increases ALLC degradation (15). This indicates that co-activation of these transcription factors integrates distinct functional aspects of independent XBP1s or ATF6 activation to influence ALLC ER quality control. Consistent with this, AP-MS shows that XBP1s and ATF6 co-activation remodels the ^FT^ALLC interactome by promoting specific changes also observed following independent transcription factor activation (**Fig. 2C,E** and **Supplemental Table 2**). For example, XBP1s and ATF6 co-activation reduces interactions between ^FT^ALLC and numerous high confidence interactors, consistent with the moderate increase in ALLC degradation observed under these conditions. Alternatively, co-activation of these transcription factors increases interactions between ^FT^ALLC and ER proteostasis factors including BiP, GRP94, ERdj3, HYOU1, and PDIA4 – all of which are also increased following ATF6 activation alone.

Comparing the functional impact of XBP1s and/or ATF6 activation on ALLC ER quality control to the changes in the interactions between ^FT^ALLC and ER proteostasis pathways provides an opportunity to identify the ER proteostasis factors likely responsible for the regulation of ALLC secretion. ATF6 activation, in both the presence or absence of XBP1s activation, reduces ALLC secretion by 50% (15). Based on our AP-MS analysis, this reduced secretion correspond to increased interactions with a specific subset of ER proteostasis factors including BiP, GRP94, HYOU1, ERdj3, and PDIA4. We have confirmed the ATF6-dependent increase in the interactions between these ER proteostasis factors and ^FT^ALLC by IP:IB (**Fig. S2G**). This suggests that these proteostasis factors are involved in dictating the selective, ATF6-dependent reduction in destabilized ALLC secretion.

### ATF6 activation increases the interactions between ER proteostasis factors and an energetically normal LC

ATF6 activation selectively reduces secretion of destabilized ALLC relative to the energetically normal LC JTO (15). Thus, we sought to define how ATF6 activation influences the interactions between JTO and ER proteostasis factors. Initially, we directly compared the interactomes of ^FT^ALLC and ^FT^JTO in vehicle-treated HEK293^DAX^ cells using our AP-MS proteomic platform (**Fig. 1A**). In order to normalize the recovery of ER proteostasis factors in these IPs, we used peptides from the λ V_c_ domain of these LCs, which is identical for both ALLC and JTO (**Fig. S1A**). This allows us to accurately monitor the differential interactions between ER proteostasis factors and specific LCs in this experiment (**Fig. S3A**). Using this approach, we identified numerous high confidence LC interacting proteins that showed increased association with the destabilized ALLC, relative to the stable JTO (**Fig. S3B** and **Supplemental Table 3**). This includes many ER proteostasis factors identified to increase association upon ATF6 activation such as BiP and GRP94, indicating that these proteins are key determinants in dictating LC ER quality control. We confirmed the increased association of select ER proteostasis factors with ALLC by IP:IB (**Fig. S3C**).

Next, we evaluated how ATF6 activation influences the interactions between ^FT^JTO and ER proteostasis factors. Since JTO is energetically more stable than ALLC, we anticipated that the increase in interactions with JTO afforded by ATF6 activation would be significantly less than that observed for ALLC. However, we found that ATF6 activation induced an identical remodeling of the ^FT^JTO interactome to that observed for ^FT^ALLC (**Fig. 2F, Fig. S3D,E** and **Supplemental Table 3**). This indicates that ATF6-dependent increases in the interactions with ER proteostasis factors occur independent of the energetic stability of the LC. Instead, our results suggest that that selective, ATF6-dependent reductions in destabilized ALLC secretion is mediated through the activity of specific ER proteostasis pathways that selectively retain the destabilized protein within the ER.

### ATF6 transcriptionally regulates ER proteostasis factors that show increased interactions with ^FT^ALLC

ATF6 activation transcriptionally regulates the expression of multiple ER proteostasis factors that show increased association with ^FT^ALLC following stress-independent ATF6 activation (e.g., BiP, GRP94) (10, 12). This suggests that the increased interaction between these ER proteostasis and ^FT^ALLC is regulated by ATF6-dependent increases in ER proteostasis factor expression. Consistent with this, ATF6-dependent changes in mRNA for high confidence interactors correlate to changes in interactions with ^FT^ALLC (**Fig. 3A** and **Supplemental Table 4**)(16). A similar relationship was observed when we compared ATF6-dependent increases in the protein levels for these ER proteostasis factors (measured by whole cell quantitative proteomics (16)) to increases in ^FT^ALLC interactions (**Fig. 3B**). These results indicate that the increased interactions between ^FT^ALLC and ER proteostasis factors is primarily dictated by ATF6-dependent increases in their expression.

**Figure 3.**
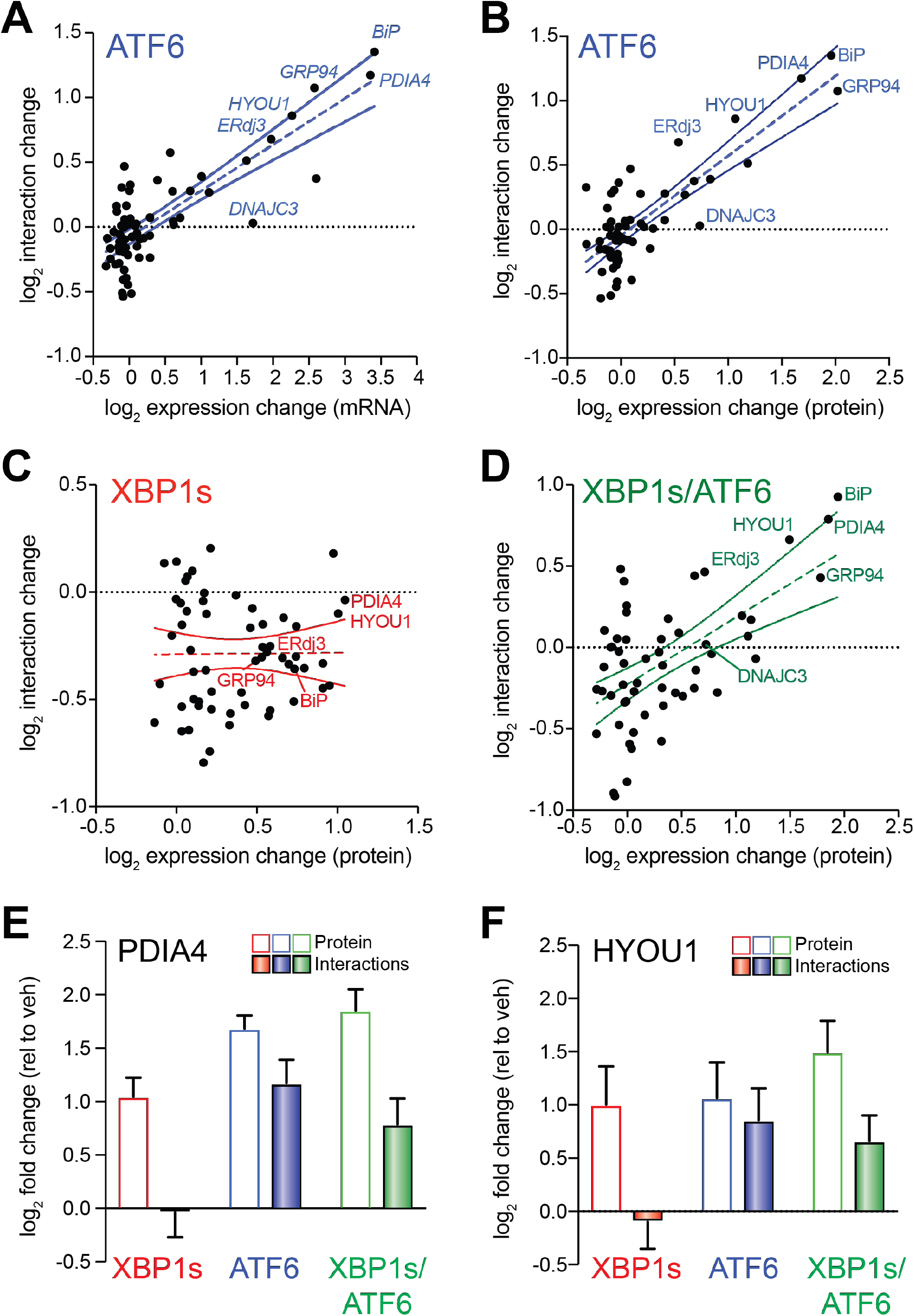
ATF6-dependent increases in ALLC interactions correlate with ER proteostasis factor expression. **A**. Plot comparing mRNA level for high confidence ALLC interacting proteins (measured by RNAseq in (16)) vs. their increased interactions with ^FT^ALLC in HEK293^DAX^ cells following stress-independent ATF6 activation. The dashed line shows least-squares linear regression. The solid lines show 95% confidence intervals. **B**. Plot comparing cellular protein level for high confidence ALLC interacting proteins (measured by whole cell quantitative proteomics in (16)) vs. their increased interactions with ^FT^ALLC in HEK293^DAX^ cells following stress-independent ATF6 activation. The dashed line shows least-squares linear regression. The solid lines show 95% confidence intervals. **C**. Plot comparing cellular protein level for high confidence ALLC interacting proteins (measured by whole cell quantitative proteomics in (16)) vs. their increased interactions with ^FT^ALLC in HEK293^DAX^ cells following stress-independent XBP1s activation. The dashed line shows least-squares linear regression. The solid lines show 95% confidence intervals. **D**. Plot comparing cellular protein level for high confidence ALLC interacting proteins (measured by whole cell quantitative proteomics in (16)) vs. their increased interactions with ^FT^ALLC in HEK293^DAX^ cells following stress-independent ATF6 and XBP1s co-activation. The dashed line shows least-squares linear regression. The solid lines show 95% confidence intervals. **E**. Graph showing changes in protein levels (open symbols) or ^FT^ALLC interactions (solid bars) for PDIA4 in HEK293^DAX^ cells following stress-independent XBP1s (red), ATF6 (blue), or XBP1s and ATF6 (green) activation. Error bars show SEM for n>3 individual replicates. **F**. Graph showing changes in protein levels (open symbols) or ^FT^ALLC interactions (solid bars) for HYOU1 in HEK293^DAX^ cells following stress-independent XBP1s (red), ATF6 (blue), or XBP1s and ATF6 (green) activation. Error bars show SEM for n>3 individual replicates.

Despite this general correlation, increased expression of ER proteostasis factors does not appear sufficient to increase ^FT^ALLC interactions. This is evident by monitoring the recovery of the high confidence LC interactor DNAJC3 in ^FT^ALLC IPs (**Fig. 3A,B**). DNAJC3 is an ER HSP40 co-chaperone that binds to misfolded proteins within the ER and directs them to the ER HSP70 BiP for ATP-dependent chaperoning (47, 48). ATF6 activation increases the expression of DNAJC3 >2-fold; however, we observe no significant increase in the association between DNAJC3 and ^FT^ALLC by AP-MS (**Fig. 3A,B** and **Fig. S4A**). This suggests that while ATF6-dependent increases in the expression of ER proteostasis factors such as BiP or GRP94 is important for dictating their increased interactions with ^FT^ALLC, increased expression does not appear sufficient to increase these interactions.

ATF6 and XBP1s induce overlapping, but distinct, subsets of ER proteostasis factors (10, 12). This provides a unique opportunity to identify key ER proteostasis factors specifically required for ATF6-dependent reductions in ALLC secretion. Towards that aim, we compared XBP1s-dependent changes in ER proteostasis factor expression to changes in their interaction with ^FT^ALLC. Unlike what we observed with ATF6 activation, ER proteostasis factor expression does not correlate with ^FT^ALLC interactions (**Fig. 3C**). However, co-activation of XBP1s and ATF6 largely restored the correlation between ER proteostasis factor expression and ^FT^ALLC interactions (**Fig. 3D**). Interestingly, specific ER proteostasis factors such as HYOU1 and PDIA4 were transcriptionally induced by XBP1s or ATF6 activation alone, but only show increased interactions with ^FT^ALLC following ATF6 activation (**Fig. 3E,F**). This is in contrast to other ER proteostasis factors such as BiP and GRP94 that are primarily regulated by ATF6 and show increased association with ^FT^ALLC following ATF6 activation (**Fig. S4B,C**). The inability for XBP1s-dependent upregulation of PDIA4 and HYOU1 to increase interactions with ^FT^ALLC suggests that the increased expression of these ER proteostasis factors is not sufficient to influence LC ER quality control. Instead, these results suggest increased targeting to ATF6-regulated, ATP-dependent chaperones such as BiP and GRP94 is primarily responsible for the ATF6-dependent increase in LC ER quality control.

### Overexpression of specific ER proteostasis factors recapitulates selective, ATF6-dependent reductions in destabilized LC secretion

Many of the ER proteostasis factors found to increase interactions with destabilized ^FT^ALLC following ATF6 activation (e.g., BiP, GRP94, and ERdj3) were previously reported to function as *‘pro-folding’* factors for LCs within the ER. BiP and GRP94 function sequentially in the folding of LCs in the ER (33). Furthermore, BiP and ERdj3 can bind multiple hydrophobic sites localized throughout a non-secreted LC, preventing its aggregation and/or premature degradation (38). In contrast, other BiP co-chaperones such as ERdj4 and ERdj5 – neither of which is regulated by ATF6 (10) – bind rarer, aggregation-prone sequences within the LC to increase its targeting to degradation. This indicates that ATF6 activation induces selective remodeling of ER chaperoning pathways that increase targeting of LCs to ATF6-regulated *‘pro-folding’* factors.

Our results indicate that ATF6 activation increases LC targeting to these *‘pro-folding’* factors by increasing their expression. Thus, we predicated that overexpression of specific *pro-folding* chaperones should mimic the capacity for ATF6 activation to selectively reduce secretion of destabilized, aggregation-prone LCs. To test this prediction, we co-overexpressed ^FT^ALLC and the ATF6-regulated chaperones BiP, GRP94, or ERdj3 in HEK293^DAX^ cells and evaluated ^FT^ALLC secretion by ELISA. In this experiment, we collected lysates and conditioned media from cells following 0 or 4 h incubation with cycloheximide (CHX) in fresh media. We then calculated fraction ^FT^ALLC secreted using the equation: fraction secreted = ^FT^ALLC media at t = 4h / ^FT^ALLC lysate at t = 0 h. Overexpression of BiP or GRP94 decreased ^FT^ALLC fraction secreted by >20%, while ERdj3 overexpression reduced ^FT^ALLC secretion by a more modest 10% (**Fig. 4A**). Similar results were observed by [^35^S] metabolic labeling (**Fig. S5A,B**). Importantly, we do not observe significant loss of ^FT^ALLC over a 4 h time course in our [^35^S] metabolic labeling experiment, indicating that the reduction in ^FT^ALLC secretion observed upon overexpression of ER chaperones does not correspond to an increase in degradation (**Fig. S5C**). This result is identical to that observed upon ATF6 activation and indicates overexpression of ER chaperones attenuate ALLC secretion through the same ER retention mechanism afforded by ATF6 activation (15).

**Figure 4.**
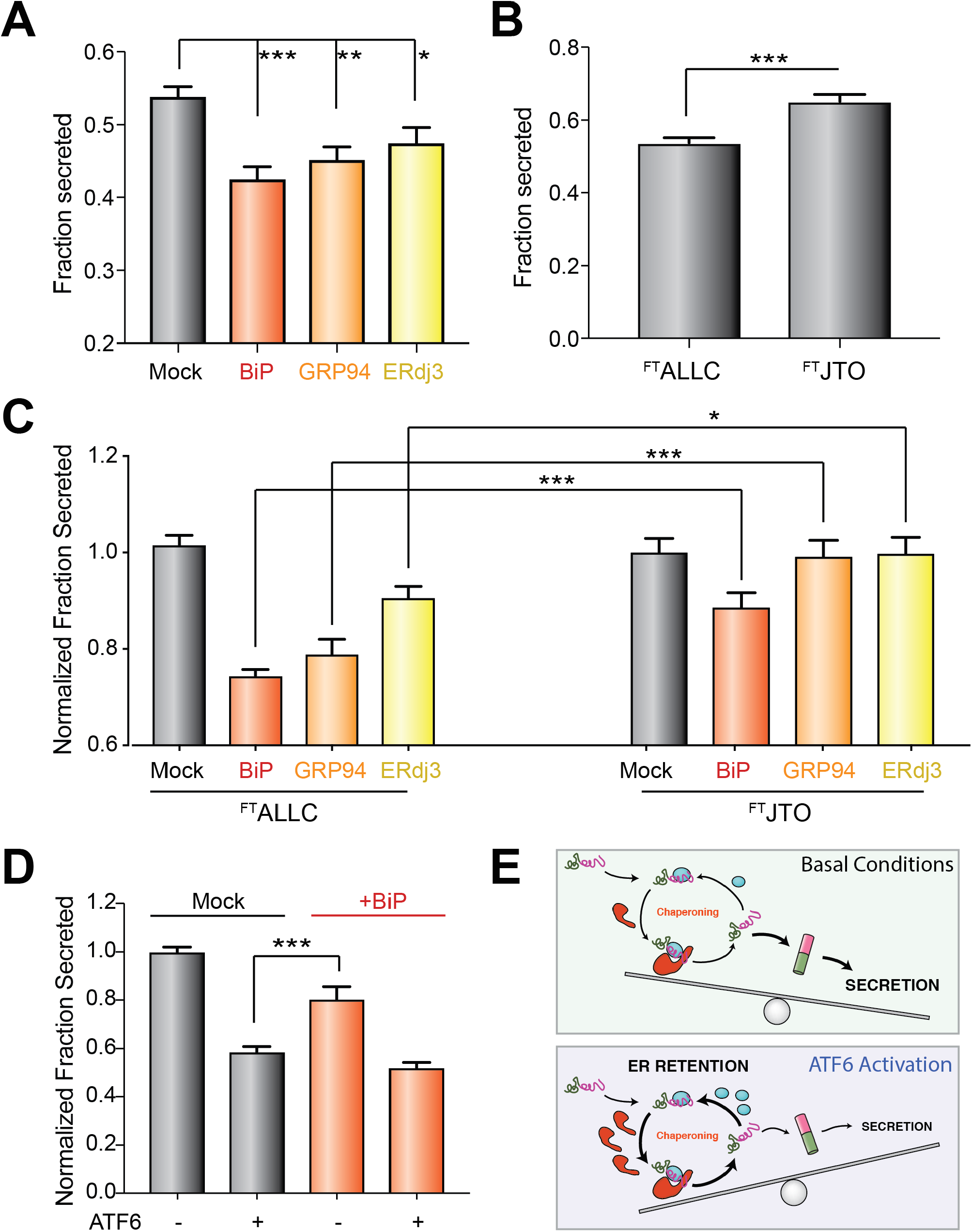
Overexpression of ATF6-regulated pro-folding ER proteostasis factors preferentially reduces ALLC secretion. **A**. Graph showing the fraction secreted of ^FT^ALLC from HEK293^DAX^ cells overexpressing the indicated ER proteostasis factor and treated with cycloheximide (CHX) for 0 or 4 h, as measured by ELISA. Fraction secreted was quantified using the following equation: fraction secreted = [^FT^ALLC in media at t=4 h] / [^FT^ALLC in lysate at t = 0 h]. Error bars show SEM for n > 14 replicates across > 4 independent experiments. *p<0.05, **p<0.01, ***p<0.005 for unpaired t-tests are shown. **B**. Graph showing fraction secretion for ^FT^ALLC or ^FT^JTO from HEK293^DAX^ cells treated with CHX for 0 or 4 h, as measured by ELISA. Fraction secreted was calculated as described in **Fig. 4A**. Error bars show SEM for n>9 replicates across n>3 independent experiments. ***p<0.005 for unpaired t-test is shown **C**. Graph showing the normalized fraction secreted of ^FT^ALLC or ^FT^JTO from HEK293^DAX^ cells overexpressing the indicated ER chaperoning factor. Normalized fraction secreted was calculated by the following equation: fraction secretion in cells overexpressing a given chaperone / fraction secretion in mock-transfected cells. Fraction secreted was calculated as in **Fig. 4A**. Error bars show SEM for n > 9 replicates collected across > 3 independent experiments. *p<0.05, ***p<0.005 for unpaired t-tests are shown. **D**. Graph showing the normalized fraction secretion of ^FT^ALLC in HEK293^DAX^ cells mock-transfected or overexpressing *BiP* subjected to a 16 h pretreatment with vehicle or ATF6 activation, as measured by ELISA. ATF6 was activated in these cells using trimethoprim (TMP; 10 μM), as previously described (10). Error bars show SEM for n=6 replicates across two independent experiments. ***p<0.005 for unpaired t-test. **E**. Illustration showing a molecular model that explains the selective, ATF6-dependent reduction in destabilized ALLC secretion. The increased chaperoning environment afforded by ATF6 activation promotes iterative rounds of ALLC chaperoning to reduce ALLC folding into a trafficking competent conformation. This leads to increased ER retention of destabilized ALLC in chaperone-bound complexes that prevent its secretion to downstream secretory environments.

ATF6 activation selectively reduces secretion of destabilized, amyloidogenic ALLC relative to the energetically-normal, non-amyloidogenic JTO. Thus, we sought to define whether overexpression of BiP, GRP94, or ERdj3 influenced secretion of ^FT^JTO using our ELISA assay (15, 16). ALLC and JTO are secreted from cells with different secretion efficiencies, reflecting differences in the ER quality control for LCs with distinct stabilities (15) – a difference further supported herein by the differential interactions between these LCs and ER proteostasis factors defined by our AP-MS analysis (**Fig. S3A-C**). Consistent with this, we found that the fraction ALLC secreted measured by CHX/ELISA is less than that observed for the more stable JTO (**Fig. 4B**). Thus, in order to compare the secretion of ALLC and JTO in cells co-overexpressing specific ER proteostasis factors, we normalized the secretion of these two LCs to control cells overexpressing each LC alone. Using this approach, we show that overexpression of BiP modestly reduces ^FT^JTO secretion; however, this reduction is significantly less than that observed for ^FT^ALLC (**Fig. 4C**). Alternatively, neither GRP94 nor ERdj3 overexpression impacted ^FT^JTO secretion. These results show that overexpression of these ER proteostasis factors preferentially reduce secretion of the destabilized, amyloidogenic ALLC, mirroring the improved LC ER quality control observed upon ATF6 activation (15).

RNAi-depletion of core ER chaperones such as BiP or GRP94 activates the UPR, preventing us from defining the importance of these ER proteostasis factors for the ATF6-dependent reduction in destabilized ALLC secretion (15). Instead, we evaluated how overexpression of the ATF6-regulated ER chaperone BiP influences ALLC secretion in the presence or absence of ATF6 activation. We selected BiP for this experiment because it is a core ER proteostasis factor whose overexpression reduces ALLC secretion to the greatest extent (**Fig. 4C**). Interestingly, the 20% reduction in ALLC secretion afforded by BiP overexpression is significantly less than the 40% reduction in secretion observed following ATF6 activation (**Fig. 4D**). Furthermore, the combination of BiP overexpression and ATF6 activation does not show significant cooperative reductions in destabilized ALLC secretion (**Fig. 4D**). Similar results were observed by [^35^S] metabolic labeling (**Fig. S5D,E**). These results indicate that overexpression of a core ER proteostasis factors only partially mimics the improved LC ER quality control observed following ATF6 activation and that maximal reductions in ALLC secretion are only achieved upon global, ATF6-dependent remodeling of ER quality control pathways.

#### Concluding Remarks

Here, we show that ATF6 activation improves ER quality control for destabilized LCs through increased targeting to select *‘pro-folding’* ER proteostasis factors. Interestingly, despite the fact that activating ATF6 selectively reduces secretion of destabilized LCs, ATF6 activation increases interactions between *‘pro-folding’* ER proteostasis factors and both destabilized (e.g., ALLC) and stable (e.g., JTO) LCs. This suggests the increased activity of these *‘pro-folding’* factors improves their capacity to *‘read-out* the energetic stability of LCs and more efficiently regulate their ER quality. A potential explanation for this effect is that increased targeting to *‘pro-folding’* ER proteostasis factors increases iterative rounds of chaperone-assisted folding that selectively prevents destabilized, amyloidogenic LCs such as ALLC from adopting a secretion-competent conformation (**Fig. 4E**). In this model, destabilized ALLC is unable to complete its folding upon release from ER chaperoning pathways. Instead, the enhanced activity of these proteostasis factors afforded by ATF6 activation promotes reengagement of ALLC prior to folding, preventing trafficking to downstream secretory environments. This reengagement of destabilized ALLC with *‘pro-folding’* factors similarly prevents targeting to degradation pathways, resulting in the ER retention observed following ATF6 activation (15). In contrast, energetically normal LCs such as JTO can efficiently fold following release from chaperoning pathways in the ATF6-remodeled ER environment due to its increased stability relative to ALLC (15, 49). This allows JTO to adopt a trafficking-competent conformation that can then be secreted to the extracellular space. Thus, while ATF6 activation increases interactions between JTO and select ER proteostasis factors, the capacity for this energetically-normal LC to fold following release from ER chaperones prevents ATF6 activation from significantly impairing its secretion. This indicates that the selective, ATF6-mediated remodeling of *‘pro-folding’* LC chaperoning pathways provides a unique opportunity to engage non-native LC conformations through interactions with multiple ER chaperones and co-chaperones to selectively reduce secretion of destabilized LCs implicated in AL disease pathogenesis.

Interestingly, overexpression of specific ER chaperones such as BiP only partially mimic the increases in LC ER quality afforded by ATF6 activation. This highlights a unique advantage for targeting endogenous transcriptional signaling pathways such as ATF6 to influence ER quality control for disease-associated proteins, as compared to targeting the activity of specific chaperones. The ATF6 transcriptional signaling pathway evolved to restore ER quality and function following diverse types of ER insults. As such, ATF6 regulates a distinct subset of ER proteostasis factors that can coordinate to impact ER quality control, providing an optimized environment to selectively influence the secretion of destabilized, amyloidogenic proteins such as amyloidogenic LCs. Consistent with this, our results show that ATF6 activation improves LC ER quality control to greater extents to that achieved by overexpression of specific ER proteostasis factors such as BiP or GRP94. This reflects the more global, ATF6-dependent remodeling of the ALLC interactome described herein, where ATF6 activation increases the interactions between ^FT^ALLC and multiple *‘pro-folding’* ER proteostasis factors.

The capacity for ATF6 activation to optimize ER proteostasis remodeling to improve LC ER quality control suggests that ATF6 activation could similarly influence the secretion of other destabilized, aggregation proteins apart from amyloidogenic LCs. Consistent with this, stress-independent ATF6 activation reduces the secretion and toxic aggregation of destabilized variants of multiple other disease-associated proteins including TTR, rhodopsin, and α1-antitrypsin (10, 17–19). Our results defining the global remodeling of ALLC interactions afforded by ATF6 activation provides a molecular basis to deconvolute the impact of ATF6 activation on the ER quality control for these and other disease-relevant proteins. Our results also further motivate the discovery of pharmacologic ATF6 activating compounds that have the potential to ameliorate the aberrant secretion and toxic aggregation of destabilized, aggregation-prone proteins implicated in etiologically-diverse protein aggregation diseases.

## MATERIALS AND METHODS

### Affinity-purification Mass Spectrometry (AP-MS) and TMT or SILAC Quantification

In general, a 10 cm tissue culture plate of HEK293^DAX^ cells was transfected with the appropriate LC expression plasmids and a fully confluent plate (approximately 10^7^ cells) was used per condition. Cell harvest, crosslinking, lysis and co-immunoprecipitation were carried out as described in the Supplemental Materials and Methods. Proteins were eluted from anti-M1 FLAG agarose beads (Sigma) twice in 75μL elution buffer (10mM Tris [pH 7.5], 2% SDS, 1mM EDTA) by heating to 95°C for 5 min. Eluted fractions were combined and proteins were precipitated in methanol/chloroform, washed twice in methanol, and then air dried. For SILAC experiments, protein pellets were resuspended in 50μL 8M urea, 50mM Tris pH 8.0, reduced with 10mM TCEP (ThermoFisher) for 30 min at room temperature, and alkylated with 12 mM iodoacetamide (Sigma) for 30min in the dark. Samples were then diluted four-fold in 50mM Tris to lower the urea concentration. For TMT experiments, the protein pellets were resuspended in 3 – 5μL 1% RapiGest SF Surfactant (Waters) followed by addition of HEPES buffer (pH 8.0, 50 mM) to a volume of 50μL. Samples were reduced with 5mM TCEP for 30min at room temperature and alkylated with 10mM iodoacetamide for 30min in the dark. Trypsin (0.5μg, Sequencing grade, Promega) was then added to the SILAC or TMT samples and incubated for 16 hours at 37°C while shaking. After digestion, SILAC peptides samples were acidified with formic acid (5% final concentration) and directly proceeded to LC-MS analysis. TMT samples were first reacted with NHS-modified TMT sixplex reagents (ThermoFisher) in 40% v/v acetonitrile and incubated for 60 min at room temperature. Reactions were then quenched by addition of 0.4% (w/v) ammonium bicarbonate. The digested and labeled samples for a given sixplex experiment were pooled and acidified with formic acid (5% final concentration). Samples were concentrated on a SpeedVac and rediluted in buffer A (94.9% water, 5% acetonitrile, 0.1 formic acid, v/v/v). Cleaved Rapigest SF and debris was removed by centrifugation for 30min at 18,000x g.

MuDPIT microcolumns were prepared as described (50), peptide samples were directly loaded onto the columns using a high-pressure chamber (Shotgun Proteomics Inc), and the columns were washed for 30min with buffer A. LC-MS/MS analysis was performed using a Q-Exactive mass spectrometer equipped with an EASY nLC 1000 (Thermo Fisher). MuDPIT experiments were performed by 10 μL sequential injections of 0, 20, 50, 80, 100% buffer C (500 mM ammonium acetate in buffer A) and a final step of 90% buffer C / 10% buffer B (19.9% water, 80% acetonitrile, 0.1% formic acid, v/v/v) and each step followed by a gradient from buffer A to buffer B on a 18 cm fused silica microcapillary column (ID 100μm) ending in a laser-pulled tip filled with Aqua C18, 3μm, 100Å resin (Phenomenex). Electrospray ionization (ESI) was performed directly from the analytical column by applying a voltage of 2.5 kV with an inlet capillary temperature of 275°C. Data-dependent acquisition of MS/MS spectra was performed with the following settings: eluted peptides were scanned from 400 to 1800 m/z with a resolution of 70,000 and the mass spectrometer in a data dependent acquisition mode. The top ten peaks for each full scan were fragmented by HCD using normalized collision energy of 30%, 2.0 m/z isolation window, 120 ms max integration time, a resolution of 7500, scanned from 100 to 1800 m/z, and dynamic exclusion set to 60s. Peptide identification and SILAC- or TMT-based protein quantification was performed as described previously using the Integrated Proteomics Pipeline Suite IP2 (Integrated Proteomics Applications, Inc.) and modules ProLuCID, DTASelect and Census (51). Data normalization was carried out manually. For SILAC experiments, the SILAC heavy/light ratios for each quantified protein were normalized to the ratio observed for ^FT^ALLC or ^FT^JTO. Each experiment represented a comparison of an experimental condition (light sample) against a common heavy reference sample (^FT^ALLC, vehicle treated). Comparisons between experimental conditions were expressed as ratios of the LC-normalized SILAC ratios. For TMT experiments, the unnormalized TMT reporter ion intensities for each quantified protein were normalized against the intensities observed for ^FT^ALLC according to the following formula: (1) 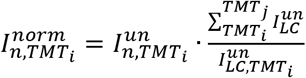, where 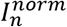 and 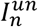 are the normalized and unnormalized TMT intensities, respectively, for a given protein *n* in the TMT channels *i-j*. Channels that did not contain LC (e.g. control transfections with untagged ALLC) were omitted from the normalization. For interactome comparison between ^FT^ALLC and ^FT^JTO, only shared peptides from the λ V_c_ constant domain were considered for the normalization. Interaction fold changes were expressed as log_2_ differences of the normalized TMT intensities for a given protein between respective TMT channels (experimental conditions), according to the following formula: (2) 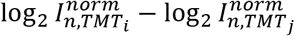. The mean of the log_2_ interaction difference was calculated from multiple MuDPIT LC-MS runs, which each represent an individual biological replicate. Significance of interaction differences was assessed by a two-tailed unpaired student’s t-test of the normalized log2-transformed TMT intensities, followed by multiple-testing correction via FDR estimation using the method of Storey et al. (52).

### Light Chain ELISA

Transfected HEK293^DAX^ were plated 150,000 cells/well in 2 identical 48-well plates (Genessee Scientific) containing 500 μL of media. Media was removed and wells were washed two times with 250 μL media containing 50 μg/mL cycloheximide (CHX). One plate was washed two times with 1x PBS and cell lysates prepared in RIPA buffer. This sample was used to monitor lysate levels of LC at t=0 h. The second plate was incubated 250 μL media with CHX for 4 h and conditioned media was collected. This sample was used to monitor secreted LC levels at 4 h. Free LC concentrations were determined by ELISA in 96-well plates (Immulon 4HBX, Thermo Fisher), as previously described (15, 16). Briefly, wells were coated overnight at 37 °C with sheep polyclonal free λ LC antibody (Bethyl Laboratories, A80-127A) at a 1:500 dilution in 50mM sodium carbonate (pH 9.6). In between all incubation steps, the plates were rinsed extensively with Tris-buffered saline containing 0.05% Tween-20 (TBST). Plates were blocked with 5% non-fat dry milk in TBST for 1 hr at 37°C. Media analytes were diluted between 5 – 200-fold in 5% non-fat dry milk in TBST and 100 μL of each sample was added to individual wells. Light chain standards ranging from 3 – 300 ng/mL were prepared from purified human Bence Jones λ light chain (Bethyl Laboratories, P80-127). Plates were incubated at 37 °C for 1.5 h with shaking. Finally, HRP-conjugated goat anti-human λ light chain antibody (Bethyl Laboratories, A80-116P) was added at a 1: 5,000 dilution in 5% non-fat dry milk in TBST, followed by a 1.5 h incubation of the plates at 37 °C. The detection was carried out with 2,2′-azinobis(3-ethylbenzothiazoline-6-sulfonic acid) (ABTS, 0.18 mg/mL) and 0.03% hydrogen peroxide in 100 mM sodium citrate pH 4.0. Detection solution (100 μL) was added to each well and the plates were incubated at room temperature. The absorbance was recorded at 405 nm and the values for the LC standards were fitted to a 4-parameter logistic function. LC concentrations were averaged from at least 3 independent replicates under each treatment and then normalized to vehicle conditions. Fraction secreted was then calculated using the equation: fraction secreted = [LC] in media at t=4 h / [LC] lysate at t = 0 h.

### Statistical Analysis

Data are presented as mean ± SEM and were analyzed by two-tailed Student’s t test to determine significance, unless otherwise indicated.

## ACKNOWLEDGEMENTS

We thank Evan Powers for critical reading of the manuscript. We thank NIH (DK107604, RLW) for financial support. LP and BR were both supported by Leukemia and Lymphoma Society postdoctoral fellowships. JCG was supported by an American Heart Association postdoctoral fellowship.

## AUTHOR CONTRIBUTIONS

RLW and JWK guided the project. LP, BR, BN, and JCG designed and carried out experiments and analyzed the data. LP, BR, and RLW prepared the figures and wrote the manuscript.

## SUPPLEMENTAL MATERIALS AND METHODS

### Plasmids and Antibodies

Plasmids expressing ^FT^ALLC, ^FT^JTO, or untagged ALLC in the pCMV1 vector were described previously (1). ^FT^BiP and ERdj3 overexpression plasmids were used as described previously (2). The GRP94 overexpression plasmid was prepared using GRP94.pDONR223 (Addgene; Cat #82130), which was recombined into pDEST40 using Gateway cloning according to the manufacturers protocol. Primary antibodies were acquired from commercial sources and used at the indicated dilutions in Antibody Buffer (50 mM Tris [pH 7.5], 150 mM NaCl supplemented with 5% BSA and 0.1% NaN_3_). Mouse monoclonal antibodies were used to detect KDEL (1:1000, Enzo Life Sciences), M2 anti-FLAG (1:500, Sigma Aldrich), Tubulin [B-5-1-2] (1:4000,Sigma), BiP/GRP-28 (1:500, Santa Cruz Biotechnology), β-actin (1:10000, Sigma Aldrich). Polyclonal rabbit antibodies were used to detect GRP94 (1:1000, GeneTex), HYOU1 (1:1000, GeneTex), ERdj3 (DNAJB11) (1:1000, ProteinTech).

### Cell Culture and Transfections

The creation and maintenance of HEK293^DAX^ cells has been described previously (3). Briefly, HEK293^DAX^ cells were cultured in high-glucose Dulbecco’s Modified Eagle’s Medium (DMEM; Corning-Cellgro) supplemented with 10% fetal bovine serum (FBS; Omega Scientific), 2 mM L-glutamine (Gibco), 100 U*mL^−1^ penicillin, and 100 μg*mL^−1^ streptomycin (Gibco). All cells were cultured under typical tissue culture conditions (37°C, 5% CO_2_). Cells were routinely tested for mycoplasma every 6 months. No further authentication of cell lines was performed by the authors. Cells were transfected using calcium phosphate precipitation, as previously described (3). All plasmids for transfection were prepared using the Qiagen Midiprep kit according to the manufacturers protocol. For SILAC experiments, the SILAC Protein Quantification kit – DMEM (ThermoFisher) was purchased. In addition to the supplied ^13^C_6_-L-Lys, the heavy media was also supplemented with ^13^C_6_,^15^N_4_-*L*-Arg (Cambridge Isotopes Laboratories, Inc.). HEK293^DAX^ cells were cultured for a minimum of 5 passages in heavy SILAC DMEM media prior to transfection.

### Immunoblotting, SDS-PAGE and Immunoprecipitation

Cell lysates were prepared as previously described in RIPA buffer (50 mM Tris, pH 7.5, 150 mM NaCl, 0.1 % SDS, 1% Triton X-100, 0.5% deoxycholate and protease inhibitor cocktail (Roche). Total protein concentration in cellular lysates was normalized using the Bio-Rad protein assay. Lysates were then denatured with 1X Laemmli buffer + 100 mM DTT and boiled before being separated by SDS-PAGE. Samples were transferred onto nitrocellulose membranes (Bio-Rad) for immunoblotting and blocked with 5% milk in Tris-buffered saline, 0.5 % Tween-20 (TBST) following incubation overnight at 4°C with primary antibodies. Membranes were washed in TBST, incubated with IR-Dye conjugated secondary antibodies and analyzed using Odyssey Infrared Imaging System (LI-COR Biosciences). Quantification was carried out with LI-COR Image Studio software. For immunoprecipitations, cells were washed with PBS and then treated with the indicated concentration Dithiobis(succinimidiyl propionate) (DSP) for 30 min at room temperature. The crosslinking reaction was quenched by addition of 100 mM Tris pH 7.5 for 15 min, then lysates were prepared in RIPA buffer. Total protein concentration in cellular lysates was normalized using Bio-Rad protein assay. Cell lysates were then subjected to preclearing with Sepharose 4B beads (Sigma) at 4 °C for 1 h with agitation. The precleared lysates were then subjected to immunoprecipitation with a M1 anti-Flag agarose resin (Sigma) at 4 °C overnight. After four washes in RIPA buffer, proteins were eluted by boiling in 6x Laemmli buffer and 100 mM DTT. Blots from IPs and inputs were probed with the primary antibodies. Membranes were then treated as described above.

### [^35^S] Metabolic Labeling

[^35^S] metabolic labeling experiments were performed as previously described (1, 3). Briefly, transfected HEK293^DAX^ were plated on poly-D-lysine coated 6-well plates and metabolically labeled in DMEM-Cys/-Met (Corning CellGro, Mediatech Inc., Manassas, VA) supplemented with glutamine, penicillin/streptomycin, dialyzed fetal bovine serum, and EasyTag EXPRESS [^35^S] Protein Labeling Mix (Perkin Elmer) for 30min. Cells were washed twice with complete media and incubated in pre-warmed DMEM for the indicated times. Media or lysates were harvested at the indicated times. Lysates were prepared in RIPA buffer (50mM Tris [pH 7.5], 150mM NaCl, 1% Triton X100, 0.5% sodium deoxycholate, 0.1% SDS) containing proteases inhibitors cocktail (Roche). FLAG-tagged LC variants were immunopurified using M1 anti-FLAG agarose beads (Sigma Aldrich) and washed four times with RIPA buffer. The immunoisolates were then eluted by boiling in 6X Laemmli buffer and separated on 12% SDS-PAGE. Gels were stained with Coomassie Blue, dried, exposed to phosphorimager plates (GE Healthcare, Pittsburgh, PA) and imaged by autoradiography using a Typhoon Trio Imager (GE Healthcare). Band intensities were quantified by densitometry in ImageQuant. Fraction secreted was calculated using the equation: fraction secreted = [extracellular [^35^S]-LC signal at t / (extracellular [^35^S]-LC signal at t=0 + intracellular [^35^S]-LC signal at t=0)]. Fraction remaining was calculated using the equation: [(extracellular [^35^S]-LC signal at t + intracellular [^35^S]-LC signal at t) / (extracellular [^35^S]-LC signal at t=0 + intracellular [^35^S]-LC signal at t=0)].

## SUPPLEMENTAL FIGURE LEGENDS

**Figure S1 (Supplement to Figure 1).**
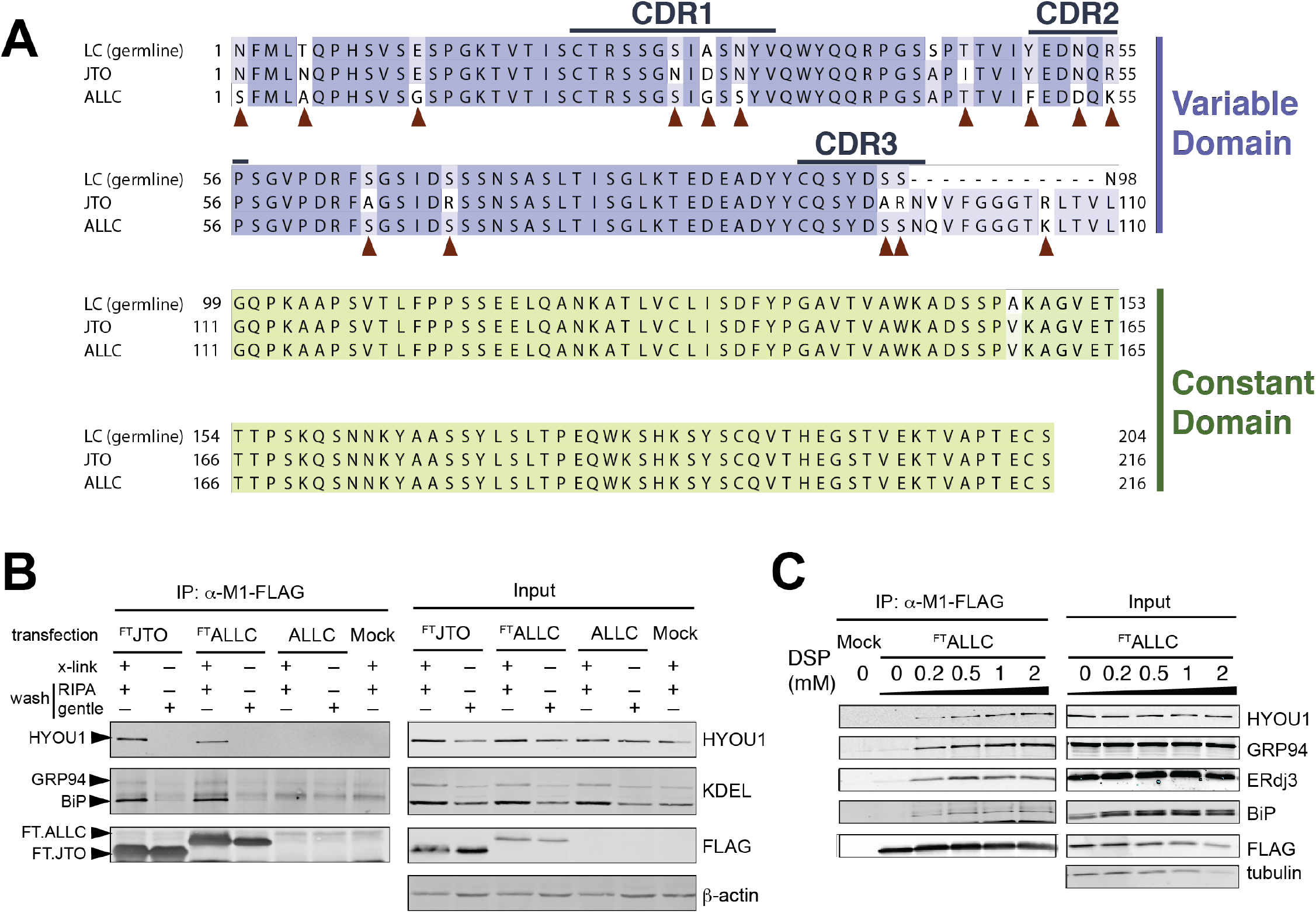
**A**. Amino acid alignment of the germline λ light chain (LC), non-amyloidogenic, energetically-normal LC JTO, and the destabilized, amyloidogenic LC ALLC used in this study. **B**. Immunoblot of Flag M1 immunopurifications (IP) prepared from HEK293^DAX^ cells transiently transfected with ^FT^JTO, ^FT^ALLC, untagged ALLC, or mock, as indicated. DSP crosslinking (0.5 mM, x-link) was added to cells prior to lysis where indicated. IPs were washed with either high-detergent RIPA or the more gentle lysis buffer (20 mM Hepes pH 7.5 100 mM NaCl 1% Triton X100), as indicated. Notice that the addition of crosslinker allows IPs to be washed with high-detergent RIPA buffer while retaining interactions with ER proteostasis buffers that are lost in the absence of crosslinking. Lysate inputs are shown as controls. **C**. Immunoblot of Flag M1 immunopurifications (IP) prepared from HEK293^DAX^ cells transiently transfected with ^FT^ALLC or mock, as indicated. Cells were crosslinked with the indicated concentration of DSP prior to lysis and IP. Lysate inputs are shown as controls.

**Figure S2 (Supplement to Figure 2).**
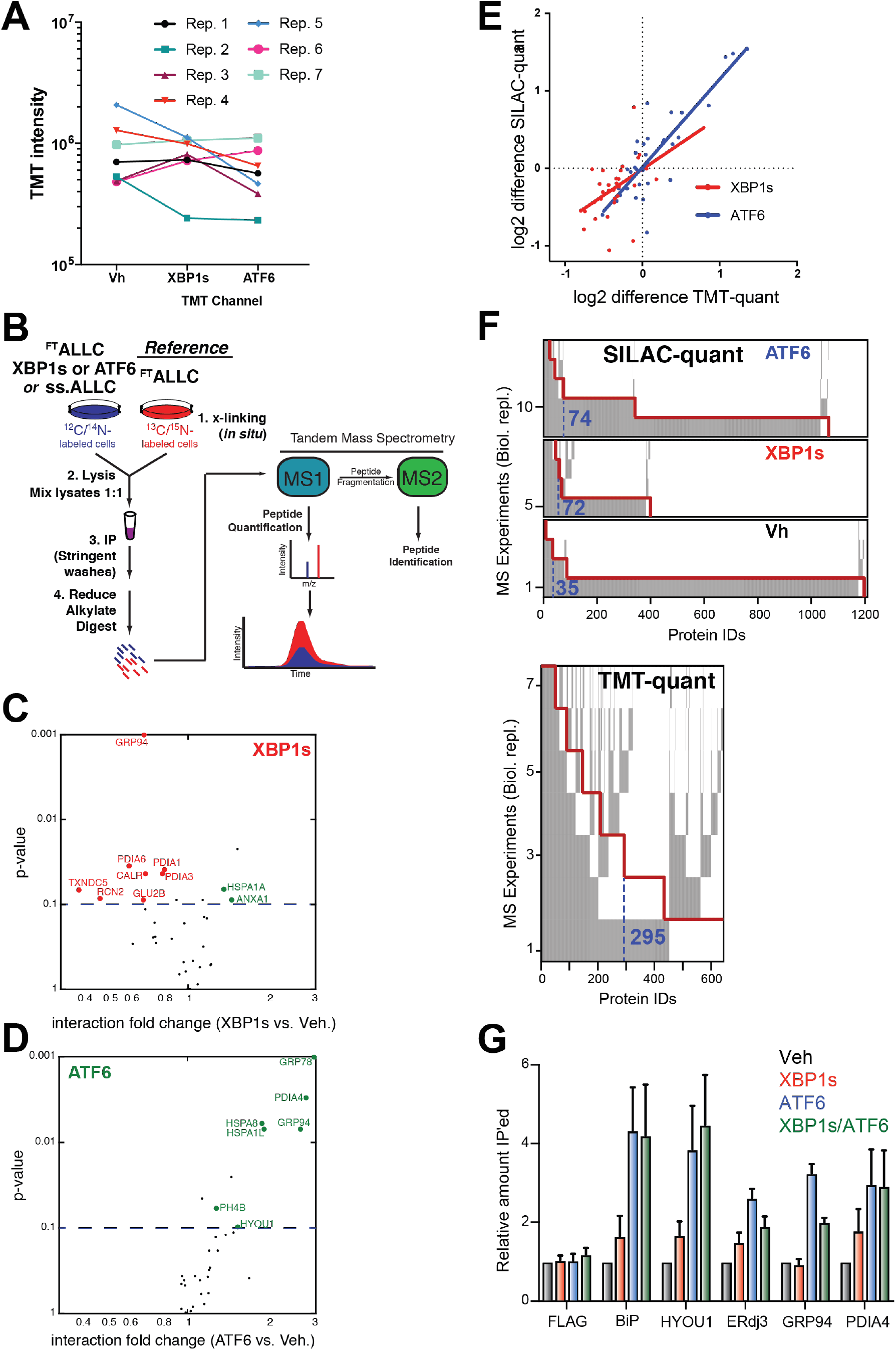
**A**. Comparison of the unnormalized mean TMT intensities for all proteins quantified in the Vh, XBP1s, and ATF6 channels of multiple replicate TMT-quant AP-MS experiments. **B**. Schematic of the SILAC-quantification based AP-MS workflow to identify interactome changes of ^FT^ALLC under conditions of stress-independent activation of XBP1s or ATF6. HEK293^DAX^ cells grown in either light ^12^C/^14^N media, or heavy ^13^C/^15^N-labeled media were transfected with ^FT^ALLC, treated with Dox or TMP to activate XBP1s or ATF6 and cross-linked in situ with DSP. Cells lysates from light and heavy cells were then mixed in equal ratios and subjected to immunoprecipitation with anti-M1 FLAG agarose beads. Protein elutions were then processed and analyzed by MuDPIT LC-MS and peptides were quantified based on the intensities of the respective heavy and light precursor ions in the MS1 chromatograms. **C,D**. Volcano plots displaying interactions changes of ^FT^ALLC measured by SILAC-quantification AP-MS after stress-independent activation of XBP1s (C) or ATF6 (D). Shown in red are negative interaction changes and in green are positive interaction changes with secretory proteins. **E**. Correlation of interactions changes observed after ATF6 (blue) or XBP1s (red) activation using TMT quantification and SILAC quantification shows good agreement. **F**. Comparison of quantified protein IDs across replicates highlights the improved detection of interaction partners using the TMT quantification approach in contrast to SILAC quantification. Highlighted in grey are proteins identified in a particular biological replicate MS experiment, and the red line shows the cumulative number of proteins quantified for the given number of replicates. For clarity of comparison, the number of proteins quantified in at least 3 replicates is listed. **G**. Graph showing relative recovery of the indicated ER proteostasis factors in ^FT^ALLC IPs following XBP1s and/or ATF6 activation in HEK293^DAX^ cells detected by immunoblotting. Error bars show SEM for n=3 replicates.

**Figure S3 (Supplement to Figure 2).**
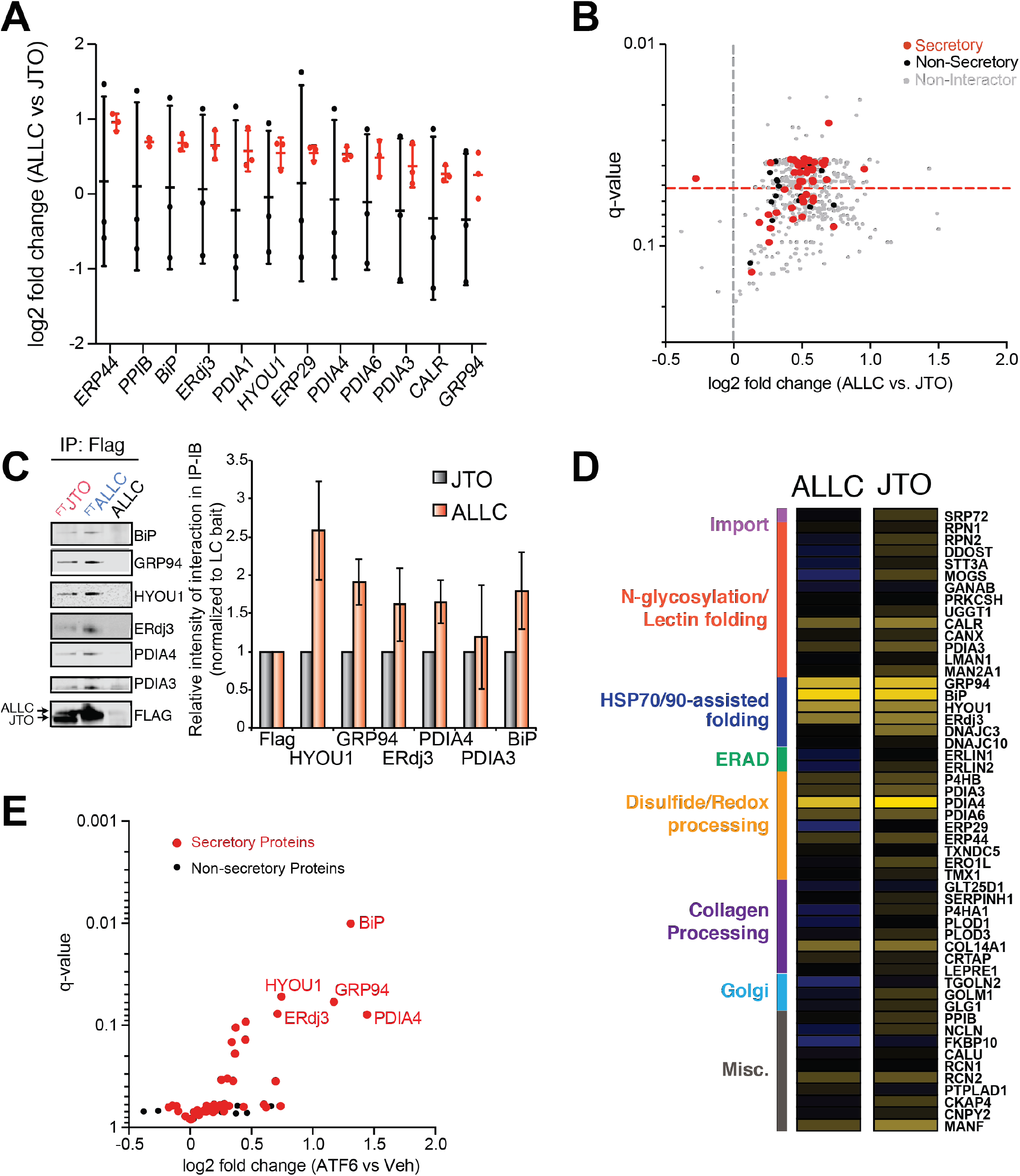
**A**. Distribution of unnormalized (black) and normalized (red) TMT ratios of ^FT^ALLC vs. ^FT^JTO for proteins with significant interaction changes. Peptides of the λ V_c_ domain, which is identical for ALLC and JTO (**Fig. S1A**), were used to normalize the TMT signal of each individual protein against the λ V_c_ domain peptide signal across each TMT channel. **B**. Plot showing TMT interaction ratio vs. q-value (Storey) for high confidence ALLC interacting proteins that co-purify with ^FT^ALLC and/or ^FT^JTO from untreated HEK293^DAX^ cells. Secretory proteins are shown in red. Full data available in **Supplemental Table 3**. **C**. Representative immunoblot of anti-FLAG IPs from HEK293^DAX^ cells transiently transfected with ^FT^JTO, ^FT^ALLC, or untagged ALLC. A graph is included showing the relative recovery of ER proteostasis factors in in ^FT^JTO (grey) or ^FT^ALLC (red) is shown. Error bars represent n = 2 independent experiments. **D**. Heatmap displaying the observed interactions changes between either ^FT^ALLC or ^FT^JTO and high confidence ER proteostasis network components following stress-independent ATF6 activation. Interactors are organized by pathway or function. **E**. Plot showing TMT interaction ratio vs. q-value for high confidence ALLC interacting proteins that co-purify with ^FT^JTO from HEK293^DAX^ cells following treatment with vehicle or TMP (to activate ATF6) for 16 h. Secretory proteins are shown in red. Full data available in **Supplemental Table 3**.

**Figure S4 (Supplement to Figure 3).**
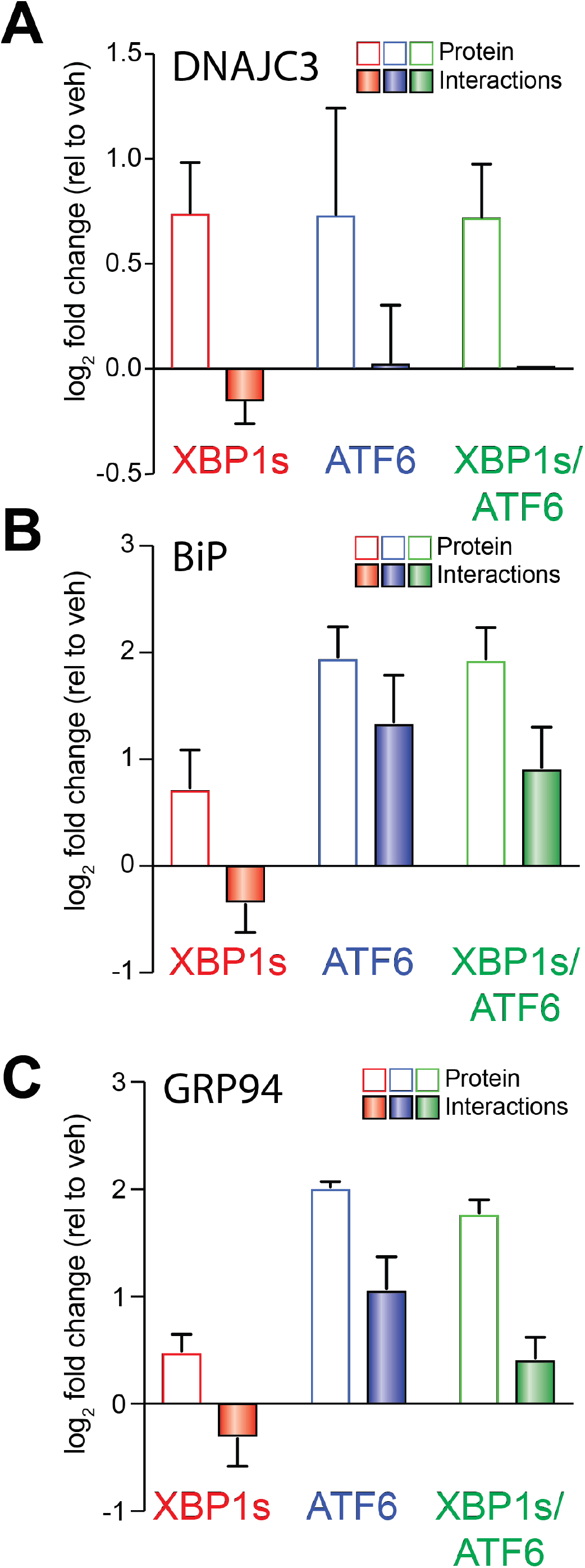
**A**. Graph showing changes in protein levels (open symbols) or ^FT^ALLC interactions (solid bars) for DNAJC3 in HEK293^DAX^ cells following stress-independent XBP1s (red), ATF6 (blue), or XBP1s and ATF6 (green) activation. Error bars show SEM for n>3 individual replicates. **B**. Graph showing changes in protein levels (open symbols) or ^FT^ALLC interactions (solid bars) for BiP in HEK293^DAX^ cells following stress-independent XBP1s (red), ATF6 (blue), or XBP1s and ATF6 (green) activation. Error bars show SEM for n>3 individual replicates. **C**. Graph showing changes in protein levels (open symbols) or ^FT^ALLC interactions (solid bars) for GRP94 in HEK293^DAX^ cells following stress-independent XBP1s (red), ATF6 (blue), or XBP1s and ATF6 (green) activation. Error bars show SEM for n>3 individual replicates.

**Figure 5 (Supplement to Figure 4).**
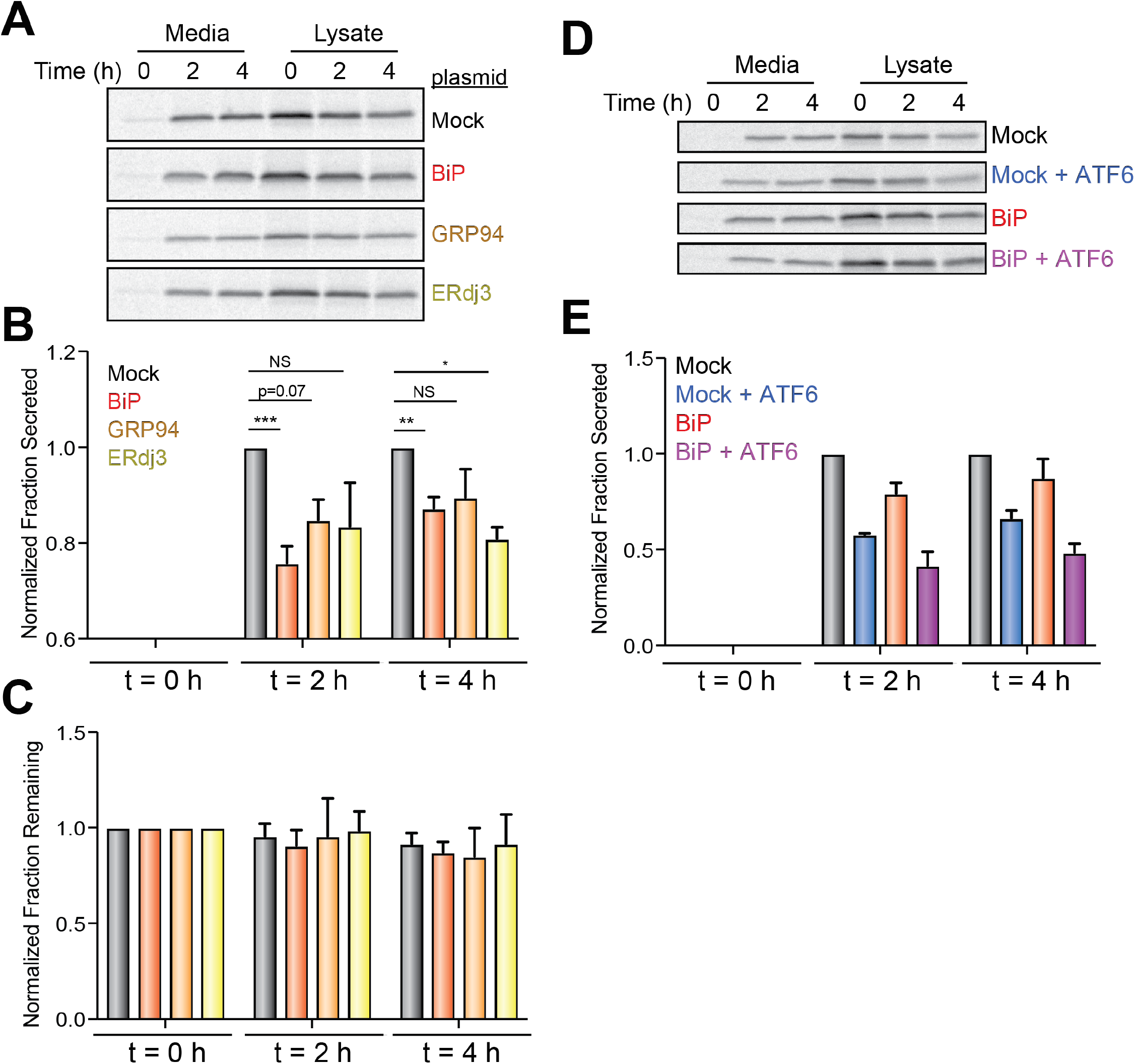
**A**. Representative autoradiogram of [^35^S]-labeled ^FT^ALLC immunopurified from lysates or media collected from HEK293^dax^ cells overexpressing mock, BiP, GRP94, or ERdj3 at the indicated time following metabolic labeling. In this experiment, cells were labeled for 30 min with [^35^S] then incubated in label free media for 0, 2 or 4 h, as described in *Supplemental Materials and Methods*. **B**. Graph showing normalized fraction [^35^S]-labeled ^FT^ALLC secreted at 0, 2 or 4 h in HEK293^DAX^ cells overexpressing mock, BiP, GRP94, or ERdj3. Fraction secreted was calculated using the following formula: fraction secreted = [^35^S]-labeled ^FT^ALLC in media at time t / ([^35^S]-labeled ^FT^ALLC in lysate at t= 0 + [^35^S]-labeled ^FT^ALLC in media at t= 0). Fraction secreted was normalized to mock transfected cells at each time point. Representative autoradiograms are shown in **Fig. S5A**. Error bars show SEM for n > 3 independent experiments. *indicates p<0.05; **indicates p<0.01; and ***indicates p<0.005 for unpaired t-tests. **C**. Graph showing fraction [^35^S]-labeled ^FT^ALLC remaining at 0, 2 or 4 h in HEK293^DAX^ cells overexpressing mock, BiP, GRP94, or ERdj3. Fraction remaining was calculated using the following formula: fraction secreted = ([^35^S]-labeled ^FT^ALLC in media at time t + [^35^S]-labeled ^FT^ALLC in lysates at time t) / ([^35^S]-labeled ^FT^ALLC in lysate at t= 0 + [^35^S]-labeled ^FT^ALLC in media at t= 0). Representative autoradiograms are shown in **Fig. S5A**. Error bars show SEM for n > 3 independent experiments. **D**. Representative autoradiogram of [^35^S]-labeled ^FT^ALLC immunopurified from lysates or media collected from HEK293^dax^ cells overexpressing mock or BiP and pretreated for 16 h with trimethoprim (TMP; 10 μM) to activate DHFR.ATF6. In this experiment, cells were labeled for 30 min with [^35^S] then incubated in label free media for 0, 2 or 4 h, as described in *Supplemental Materials and Methods*. **E**. Graph showing normalized fraction [^35^S]-labeled ^FT^ALLC secreted at 0, 2 or 4 h in HEK293^DAX^ cells overexpressing mock or BiP and pretreated for 16 h with trimethoprim (TMP; 10 μM) to activate DHFR.ATF6 in these cells. Fraction secreted was calculated as described in **Fig. S5B**. Fraction secreted was normalized to mock transfected cells at each time point. Representative autoradiograms are shown in **Fig. S5D**. Error bars show SEM for n=2 independent experiments.

## SUPPLEMENTAL TABLE LEGENDS

**Supplemental Table 1 (Supplement to Figure 1)**. Excel spreadsheets including the interactome data comparing the interactions between ER proteostasis factors and either ^FT^LC (combined replicates of ^FT^ALLC and ^FT^JTO) or untagged ALLC. Two sheets are included within this file: 1) a summary sheet including only the final TMT ratios and significance and 2) a sheet containing all of the raw data for the included analyses.

**Supplemental Table 2 (Supplement to Figure 2)**. Excel spreadsheets describing the interactome data comparing interactions between ER proteostasis factors and ^FT^ALLC following stress-independent XBP1s and/or ATF6 activation in HEK293^DAX^ cells. Two sheets are included within this file: 1) a summary sheet including only the final TMT ratios and significance and 2) a sheet containing all of the raw data for the included analyses.

**Supplemental Table 3 (Supplement to Figure 2)**. Excel spreadsheet describing the interactome data comparing the interactions between ER proteostasis factors and ^FT^ALLC and ^FT^JTO in HEK293^DAX^ cells or ^FT^JTO in HEK293^DAX^ cells following stress-independent ATF6 activation. Four sheets are included within this file: 1) a summary sheet including only the final TMT ratios and significance comparing the interaction ratios between ^FT^ALLC and ^FT^JTO and 2) a sheet containing all of the raw data used to compare the interactomes of ^ft^ALLC and ^FT^JTO, 3) a summary sheet including only the final TMT ratios and significance comparing the interaction ratios for ^FT^JTO in the absence or presence of ATF6 activation in HEK293^DAX^ cells and 4) a sheet containing all of the raw data used to compare the interactome ^FT^JTO in the presence or absence of ATF6 activation.

**Supplemental Table 4 (Supplement to Figure 3)**. Excel spreadsheets comparing changes in the mRNA or protein levels and ^FT^ALLC interactions for high confidence LC interacting proteins in HEK293^DAX^ cells following stress-independent activation of ATF6, XBP1s, or ATF6 and XBP1s co-activation. This table contains four sheets. Data for changes in mRNA or protein levels in HEK293^DAX^ cells following these treatments is from (3, 4).

## REFERENCES

1. Blancas-Mejia LM, Ramirez-Alvarado M. Systemic amyloidoses. Annu Rev Biochem. 2013;82:745–74.

2. Powers ET, Morimoto RI, Dillin A, Kelly JW, Balch WE. Biological and chemical approaches to diseases of proteostasis deficiency. Annu Rev Biochem. 2009;78:959–91.

3. Plate L, Wiseman RL. Regulating Secretory Proteostasis through the Unfolded Protein Response: From Function to Therapy. Trends Cell Biol. 2017;27(10):722–37.

4. Braakman I, Bulleid NJ. Protein folding and modification in the mammalian endoplasmic reticulum. Annu Rev Biochem. 2011;80:71–99.

5. Guerriero CJ, Brodsky JL. The delicate balance between secreted protein folding and endoplasmic reticulum-associated degradation in human physiology. Physiol Rev. 2012;92(2):537–76.

6. Kim YE, Hipp MS, Bracher A, Hayer-Hartl M, Hartl FU. Molecular chaperone functions in protein folding and proteostasis. Annu Rev Biochem. 2013;82:323–55.

7. Feige MJ, Buchner J. Principles and engineering of antibody folding and assembly. Biochim Biophys Acta. 2014;1844(11):2024–31.

8. Balchin D, Hayer-Hartl M, Hartl FU. In vivo aspects of protein folding and quality control. Science. 2016;353(6294):aac4354.

9. Hetz C, Saxena S. ER stress and the unfolded protein response in neurodegeneration. Nat Rev Neurol. 2017;13(8):477–91.

10. Shoulders MD, Ryno LM, Genereux JC, Moresco JJ, Tu PG, Wu C, Yates JR, 3rd, Su AI, Kelly JW, Wiseman RL. Stress-independent activation of XBP1s and/or ATF6 reveals three functionally diverse ER proteostasis environments. Cell Rep. 2013;3(4):1279–92.

11. Lee AH, Iwakoshi NN, Glimcher LH. XBP-1 regulates a subset of endoplasmic reticulum resident chaperone genes in the unfolded protein response. Mol Cell Biol. 2003;23(21):7448–59.

12. Yamamoto K, Yoshida H, Kokame K, Kaufman RJ, Mori K. Differential contributions of ATF6 and XBP1 to the activation of endoplasmic reticulum stress-responsive cis-acting elements ERSE, UPRE and ERSE-II. J Biochem. 2004;136(3):343–50.

13. Adachi Y, Yamamoto K, Okada T, Yoshida H, Harada A, Mori K. ATF6 is a transcription factor specializing in the regulation of quality control proteins in the endoplasmic reticulum. Cell Struct Funct. 2008;33(1):75–89.

14. Arendt BK, Ramirez-Alvarado M, Sikkink LA, Keats JJ, Ahmann GJ, Dispenzieri A, Fonseca R, Ketterling RP, Knudson RA, Mulvihill EM, Tschumper RC, Wu X, Zeldenrust SR, Jelinek DF. Biologic and genetic characterization of the novel amyloidogenic lambda light chain-secreting human cell lines, ALMC-1 and ALMC-2. Blood. 2008;112(5):1931–41.

15. Cooley CB, Ryno LM, Plate L, Morgan GJ, Hulleman JD, Kelly JW, Wiseman RL. Unfolded protein response activation reduces secretion and extracellular aggregation of amyloidogenic immunoglobulin light chain. Proc Natl Acad Sci U S A. 2014;111(36):13046–51.

16. Plate L, Cooley CB, Chen JJ, Paxman RJ, Gallagher CM, Madoux F, Genereux JC, Dobbs W, Garza D, Spicer TP, Scampavia L, Brown SJ, Rosen H, Powers ET, Walter P, Hodder P, Wiseman RL, Kelly JW. Small molecule proteostasis regulators that reprogram the ER to reduce extracellular protein aggregation. Elife. 2016;5.

17. Chen JJ, Genereux JC, Qu S, Hulleman JD, Shoulders MD, Wiseman RL. ATF6 activation reduces the secretion and extracellular aggregation of destabilized variants of an amyloidogenic protein. Chem Biol. 2014;21(11):1564–74.

18. Smith SE, Granell S, Salcedo-Sicilia L, Baldini G, Egea G, Teckman JH, Baldini G. Activating transcription factor 6 limits intracellular accumulation of mutant alpha(1)-antitrypsin Z and mitochondrial damage in hepatoma cells. J Biol Chem. 2011;286(48):41563–77.

19. Chiang WC, Hiramatsu N, Messah C, Kroeger H, Lin JH. Selective activation of ATF6 and PERK endoplasmic reticulum stress signaling pathways prevent mutant rhodopsin accumulation. Invest Ophthalmol Vis Sci. 2012;53(11):7159–66.

20. Wiseman RL, Powers ET, Buxbaum JN, Kelly JW, Balch WE. An adaptable standard for protein export from the endoplasmic reticulum. Cell. 2007;131(4):809–21.

21. Kean MJ, Couzens AL, Gingras AC. Mass spectrometry approaches to study mammalian kinase and phosphatase associated proteins. Methods. 2012;57(4):400–8.

22. Pankow S, Bamberger C, Calzolari D, Bamberger A, Yates JR, 3rd. Deep interactome profiling of membrane proteins by co-interacting protein identification technology. Nat Protoc. 2016;11(12):2515–28.

23. Pankow S, Bamberger C, Calzolari D, Martinez-Bartolome S, Lavallee-Adam M, Balch WE, Yates JR 3rd., F508 CFTR interactome remodelling promotes rescue of cystic fibrosis. Nature. 2015;528(7583):510–6.

24. Taipale M, Tucker G, Peng J, Krykbaeva I, Lin ZY, Larsen B, Choi H, Berger B, Gingras AC, Lindquist S. A quantitative chaperone interaction network reveals the architecture of cellular protein homeostasis pathways. Cell. 2014;158(2):434–48.

25. Budayeva HG, Cristea IM. A mass spectrometry view of stable and transient protein interactions. Adv Exp Med Biol. 2014;806:263–82.

26. Miteva YV, Budayeva HG, Cristea IM. Proteomics-based methods for discovery, quantification, and validation of protein-protein interactions. Anal Chem. 2013;85(2):749–68.

27. Wall J, Schell M, Murphy C, Hrncic R, Stevens FJ, Solomon A. Thermodynamic instability of human lambda 6 light chains: correlation with fibrillogenicity. Biochemistry. 1999;38(42):14101–8.

28. Lomant AJ, Fairbanks G. Chemical probes of extended biological structures: synthesis and properties of the cleavable protein cross-linking reagent [35S]dithiobis(succinimidyl propionate). J Mol Biol. 1976;104(1):243–61.

29. Nittis T, Guittat L, LeDuc RD, Dao B, Duxin JP, Rohrs H, Townsend RR, Stewart SA. Revealing novel telomere proteins using in vivo cross-linking, tandem affinity purification, and label-free quantitative LC-FTICRMS. Mol Cell Proteomics. 2010;9(6):1144–56.

30. Smith AL, Friedman DB, Yu H, Carnahan RH, Reynolds AB. ReCLIP (reversible cross-link immuno-precipitation): an efficient method for interrogation of labile protein complexes. PLoS One. 2011;6(1):e16206.

31. Washburn MP, Wolters D, Yates 3rd., Large-scale analysis of the yeast proteome by multidimensional protein identification technology. Nat Biotechnol. 2001;19(3):242–7.

32. Yates JR, Ruse CI, Nakorchevsky A. Proteomics by mass spectrometry: approaches, advances, and applications. Annu Rev Biomed Eng. 2009;11:49–79.

33. Melnick J, Dul JL, Argon Y. Sequential interaction of the chaperones BiP and GRP94 with immunoglobulin chains in the endoplasmic reticulum. Nature. 1994;370(6488):373–5.

34. Davis DP, Khurana R, Meredith S, Stevens FJ, Argon Y. Mapping the major interaction between binding protein and Ig light chains to sites within the variable domain. J Immunol. 1999;163(7):3842–50.

35. Skowronek MH, Hendershot LM, Haas IG. The variable domain of nonassembled Ig light chains determines both their half-life and binding to the chaperone BiP. Proc Natl Acad Sci U S A. 1998;95(4):1574–8.

36. Hellman R, Vanhove M, Lejeune A, Stevens FJ, Hendershot LM. The in vivo association of BiP with newly synthesized proteins is dependent on the rate and stability of folding and not simply on the presence of sequences that can bind to BiP. J Cell Biol. 1999;144(1):21–30.

37. Melnick J, Aviel S, Argon Y. The endoplasmic reticulum stress protein GRP94, in addition to BiP, associates with unassembled immunoglobulin chains. J Biol Chem. 1992;267(30):21303–6.

38. Behnke J, Mann MJ, Scruggs FL, Feige MJ, Hendershot LM. Members of the Hsp70 Family Recognize Distinct Types of Sequences to Execute ER Quality Control. Mol Cell. 2016;63(5):739–52.

39. Shen Y, Hendershot LM. ERdj3, a stress-inducible endoplasmic reticulum DnaJ homologue, serves as a cofactor for BiP's interactions with unfolded substrates. Mol Biol Cell. 2005;16(1):40–50.

40. Behnke J, Feige MJ, Hendershot LM. BiP and its nucleotide exchange factors Grp170 and Sil1: mechanisms of action and biological functions. J Mol Biol. 2015;427(7):1589–608.

41. Cole KS, Grandjean JMD, Chen K, Witt CH, O’Day J, Shoulders MD, Wiseman RL, Weerapana E. Characterization of an A-Site Selective Protein Disulfide Isomerase A1 Inhibitor. Biochemistry. 2018;57(13):2035–43.

42. Hsu TA, Watson S, Eiden JJ, Betenbaugh MJ. Rescue of immunoglobulins from insolubility is facilitated by PDI in the baculovirus expression system. Protein Expr Purif. 1996;7(3):281–8.

43. Selbach M, Mann M. Protein interaction screening by quantitative immunoprecipitation combined with knockdown (QUICK). Nat Methods. 2006;3(12):981–3.

44. Ong SE, Blagoev B, Kratchmarova I, Kristensen DB, Steen H, Pandey A, Mann M. Stable isotope labeling by amino acids in cell culture, SILAC, as a simple and accurate approach to expression proteomics. Mol Cell Proteomics. 2002;1(5):376–86.

45. Kaake RM, Wang X, Huang L. Profiling of protein interaction networks of protein complexes using affinity purification and quantitative mass spectrometry. Mol Cell Proteomics. 2010;9(8):1650–65.

46. Christianson JC, Olzmann JA, Shaler TA, Sowa ME, Bennett EJ, Richter CM, Tyler RE, Greenblatt EJ, Harper JW, Kopito RR. Defining human ERAD networks through an integrative mapping strategy. Nat Cell Biol. 2011;14(1):93–105.

47. Petrova K, Oyadomari S, Hendershot LM, Ron D. Regulated association of misfolded endoplasmic reticulum lumenal proteins with P58/DNAJc3. EMBO J. 2008;27(21):2862–72.

48. Rutkowski DT, Kang SW, Goodman AG, Garrison JL, Taunton J, Katze MG, Kaufman RJ, Hegde RS. The role of p58IPK in protecting the stressed endoplasmic reticulum. Mol Biol Cell. 2007;18(9):3681–91.

49. Morgan GJ, Kelly JW. The Kinetic Stability of a Full-Length Antibody Light Chain Dimer Determines whether Endoproteolysis Can Release Amyloidogenic Variable Domains. J Mol Biol. 2016;428(21):4280–97.

50. Fonslow BR, Niessen SM, Singh M, Wong CC, Xu T, Carvalho PC, Choi J, Park SK, Yates JR, 3rd. Single-step inline hydroxyapatite enrichment facilitates identification and quantitation of phosphopeptides from mass-limited proteomes with MudPIT. J Proteome Res. 2012;11(5):2697–709.

51. Ryno LM, Genereux JC, Naito T, Morimoto RI, Powers ET, Shoulders MD, Wiseman RL. Characterizing the altered cellular proteome induced by the stress-independent activation of heat shock factor 1. ACS Chem Biol. 2014;9(6):1273–83.

52. Storey JD, Tibshirani R. Statistical significance for genomewide studies. Proc Natl Acad Sci U S A. 2003;100(16):9440–5.

## SUPPLEMENTAL REFERENCES

1. Cooley CB, Ryno LM, Plate L, Morgan GJ, Hulleman JD, Kelly JW, Wiseman RL. Unfolded protein response activation reduces secretion and extracellular aggregation of amyloidogenic immunoglobulin light chain. Proc Natl Acad Sci U S A. 2014;111(36):13046–51.

2. Genereux JC, Qu S, Zhou M, Ryno LM, Wang S, Shoulders MD, Kaufman RJ, Lasmezas CI, Kelly JW, Wiseman RL. Unfolded protein response-induced ERdj3 secretion links ER stress to extracellular proteostasis. EMBO J. 2015;34(1):4–19.

3. Shoulders MD, Ryno LM, Genereux JC, Moresco JJ, Tu PG, Wu C, Yates JR, 3rd, Su AI, Kelly JW, Wiseman RL. Stress-independent activation of XBP1s and/or ATF6 reveals three functionally diverse ER proteostasis environments. Cell Rep. 2013;3(4):1279–92.

4. Plate L, Cooley CB, Chen JJ, Paxman RJ, Gallagher CM, Madoux F, Genereux JC, Dobbs W, Garza D, Spicer TP, Scampavia L, Brown SJ, Rosen H, Powers ET, Walter P, Hodder P, Wiseman RL, Kelly JW. Small molecule proteostasis regulators that reprogram the ER to reduce extracellular protein aggregation. Elife. 2016;5.

